# Apobec-Mediated Retroviral Hypermutation *In Vivo* is Dependent on Mouse Strain

**DOI:** 10.1101/2023.11.02.565355

**Authors:** Hyewon Byun, Gurvani B. Singh, Wendy Kaichun Xu, Poulami Das, Alejandro Reyes, Anna Battenhouse, Dennis C. Wylie, Mary M. Lozano, Jaquelin P. Dudley

**Author notes:** Corresponding author: Jaquelin Dudley, Dept. of Molecular Biosciences, The University of Texas at Austin, 100 E. 24th Street, NHB 2.616, Stop 5000; Austin, TX 78712.

## Abstract

Replication of the complex retrovirus mouse mammary tumor virus (MMTV) is antagonized by murine Apobec3 (mA3), a member of the Apobec family of cytidine deaminases. We have shown that MMTV-encoded Rem protein inhibits proviral mutagenesis by the Apobec enzyme, activation-induced cytidine deaminase (AID) during viral replication in BALB/c mice. To further study the role of Rem *in vivo*, we have infected C57BL/6 (B6) mice with a superantigen-independent lymphomagenic strain of MMTV (TBLV-WT) or a mutant strain (TBLV-SD) that is defective in Rem and its cleavage product Rem-CT. Unlike MMTV, TBLV induced T-cell tumors in µMT mice, indicating that mature B cells, which express the highest AID levels, are not required for TBLV replication. Compared to BALB/c, B6 mice were more susceptible to TBLV infection and tumorigenesis. The lack of Rem expression accelerated B6 tumorigenesis at limiting doses compared to TBLV-WT in either wild-type B6 or AID-deficient mice. However, unlike proviruses from BALB/c mice, high-throughput sequencing indicated that proviral G-to-A or C-to-T changes did not significantly differ in the presence and absence of Rem expression. *Ex vivo* stimulation showed higher levels of mA3 relative to AID in B6 compared to BALB/c splenocytes, but effects of agonists differed in the two strains. RNA-Seq revealed increased transcripts related to growth factor and cytokine signaling in TBLV-SD-induced tumors relative to those from TBLV-WT, consistent with a third Rem function. Thus, Rem-mediated effects on tumorigenesis in B6 mice are independent of Apobec-mediated proviral hypermutation.

---

Retroviruses are small RNA-containing viruses that replicate through a DNA intermediate to establish lifelong infections of their hosts (1). To maintain long-term infections, retroviruses have developed various strategies to thwart the host immune response (2, 3). The human APOBEC family cytidine deaminases contribute to innate immunity against RNA- and DNA-containing viruses, including retroviruses, as well as retrotransposons (2, 4, 5). Retroviral replication often is inhibited by incorporation of these enzymes into virions (6–8) since APOBEC3 (A3) packaging results in blocks to reverse transcription and mutagenesis of the proviral genome (9, 10). A3-mediated cytidine deamination on the human immunodeficiency virus type 1 (HIV-1) DNA negative strand in the absence of Vif leads to G-to-A sequence changes on the plus strand and inhibition of replication (11–13). HIV-1-encoded Vif acts as an adapter to Cullin-based E3 ligases, which lead to ubiquitylation and proteasomal degradation of human APOBEC3G and 3F (14, 15). Nonetheless, many aspects of the immune response cannot feasibly be studied in humans. In contrast, mouse genetics has provided many important insights into the conserved biology of viruses and the antiviral immune response, including the role of Apobec proteins (16).

Experiments using the betaretrovirus mouse mammary tumor virus (MMTV) were the first to show inhibition of retroviral replication by mouse Apobec enzymes *in vivo* (6). Unlike humans, mice have a single *Apobec3* (*mA3*) gene (2). MMTV-RIII strain infection of C57BL/6 (B6) mice lacking a functional *mA3* gene showed accelerated viral replication and higher proviral loads compared to those in wild-type B6 mice. Interestingly, MMTV hypermutation was not observed in the presence of mA3 despite its packaging into MMTV virions (6). Subsequent studies indicated that mA3 blocked MMTV reverse transcription (17), suggesting that mA3 does not induce hypermutation of the MMTV proviral genome.

Our studies in BALB/c mice showed that the Apobec family enzyme activation-associated cytidine deaminase (AID) leads to MMTV hypermutation in the absence of the virally encoded Rem protein (18). MMTV proviruses lacking Rem expression (MMTV-SD) had increased cytidine mutations within the AID-associated WR<u>C</u> (W = A/T, R = A/G) sequence motif, but also in TY<u>C</u> (Y = C/T) motifs associated with mA3 expression compared to wild-type MMTV proviruses (MMTV-WT). Both types of mutations were greatly reduced in MMTV-SD-induced tumors from mice lacking AID expression (18). Transfection experiments showed that Rem co-expression leads to proteasomal degradation of AID, but not mA3. These experiments suggested that MMTV Rem is the functional equivalent of HIV-1 Vif and antagonizes AID, but possibly other Apobec family enzymes (18).

Rem is translated from a doubly spliced version of the envelope gene and in the same reading frame (19, 20). Both Env and Rem are translated at the endoplasmic reticulum (ER) membrane. Their common signal peptide (SP) is cleaved by signal peptide peptidase (21–23) and retrotranslocated to yield a Rev-like protein that traffics to the nucleus for viral RNA export and expression (21, 24, 25) (Fig. 1A). Since the MMTV-SD mutant produces SP from signal peptidase cleavage of the Env precursor, the Apobec-induced hypermutation phenotype of this mutant is due to Rem C-terminal sequences, either uncleaved Rem or the C-terminal cleavage product, Rem-CT (18). However, Rem-CT is produced entirely within the ER and traffics within endosomal membranes, suggesting that uncleaved Rem is the Apobec antagonist (26).

**Fig. 1.**
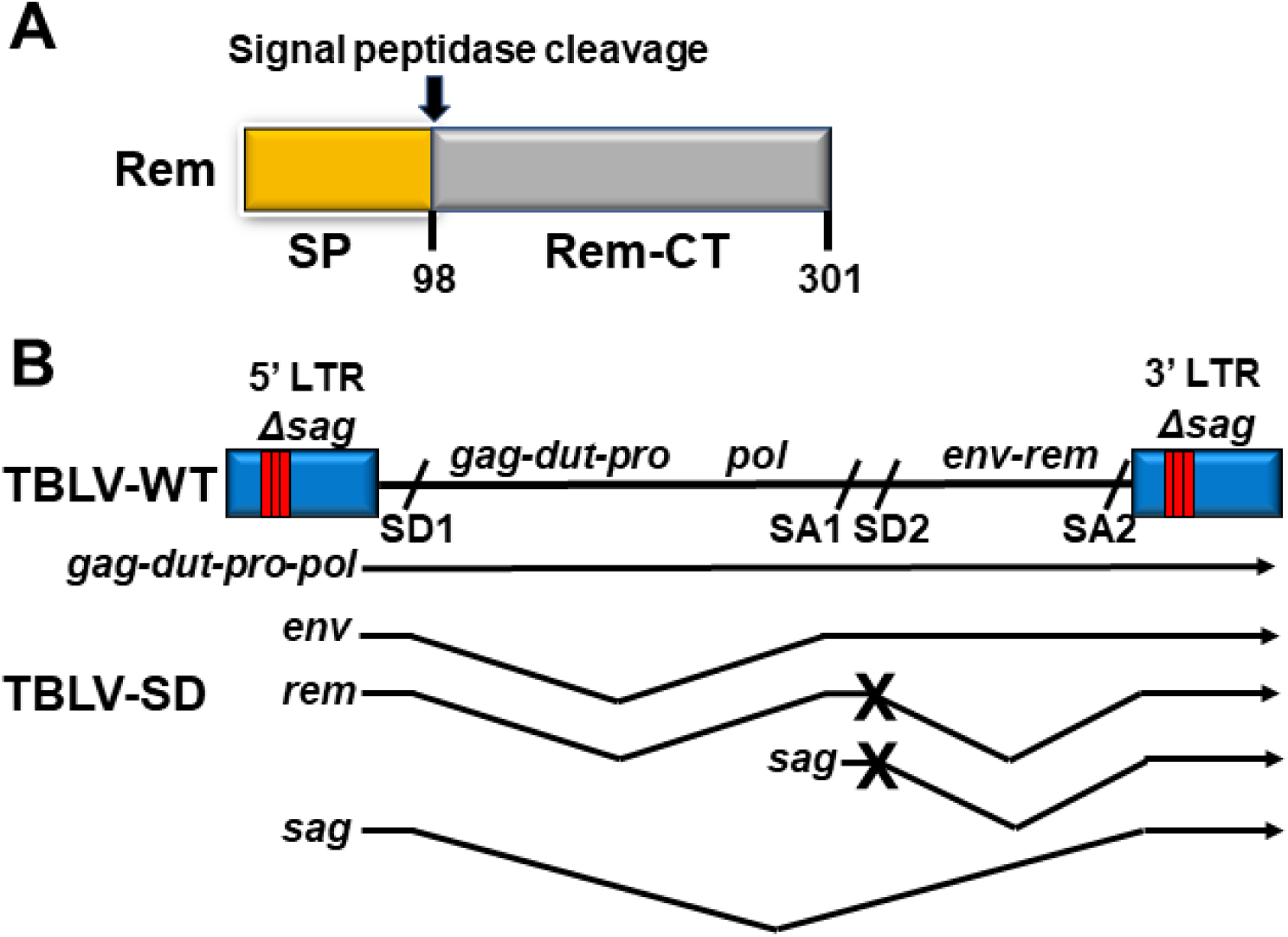
Diagram of Rem and clonal infectious TBLV proviruses. (A) Domain organization of Rem and its cleavage products SP and Rem-CT. The arrow indicates the position of Rem cleavage by signal peptidase. (B) Diagram of infectious TBLV-WT and TBLV-SD proviruses and their mRNA transcripts. The relative positions of genes are indicated on the proviral DNA. Both TBLV-WT and TBLV-SD have a T-cell enhancer that consists of a deletion and triplication of flanking sequences within the LTRs (red bars) compared to MMTV (36, 52). The LTR alteration leads to elimination of Sag expression, which is required for MMTV transmission and mammary tumorigenesis, but allows development of T-cell tumors (38). The splice donor (SD) and acceptor (SA) sites are indicated on the provirus. Transcripts are shown below TBLV-WT (arrows), and introns are shown by V shapes. The TBLV-SD provirus has a mutation (marked by an X) in SD site 2 (SD2), which prevents the synthesis of the doubly spliced *rem* mRNA (79) as well as *sag* transcripts from the intragenic promoter. Since TBLV does not have a functional *sag* open reading frame, TBLV-WT and TBLV-SD differ only by the production of *rem* mRNA.

MMTV transmission to the mammary gland prior to tumor induction requires replication in both B and T cells (27–30), which are known to express AID and mA3 (31–33). Apobec effects on retroviral replication *in vivo* primarily have been studied on the C57BL/6 (B6) background because of their ease of genetic manipulation (34), yet different mouse strains have distinct immune responses to pathogens (35). To further understand the involvement of cytidine deaminases in MMTV replication, we conducted additional experiments in B6 mice using the Sag-independent MMTV strain (TBLV-WT), which induces T-cell lymphomas rather than breast cancers (36–38) (Fig. 1B). MMTV-encoded Sag protein on the surface of mature B cells is required for MMTV transmission from maternal milk in the gut to the mammary glands (39, 40). In this report, mutant TBLV lacking Rem expression (TBLV-SD) accelerated T-cell tumor induction compared to TBLV-WT in wild-type as well as AID-deficient *Aicda*^-/-^ B6 mice (41). Unlike the increased proviral mutations observed in the absence of Rem in BALB/c mice, TBLV-WT and TBLV-SD proviruses from B6 tumors had similar frequencies of mutations in the envelope region. Furthermore, RNA-seq analysis on B6 tumors indicated that most transcriptional changes were observed in tumors from *Aicda*^-/-^ mice in the absence of Rem, including elevation of genes involved in growth factor and cytokine signaling. The results suggest that the latency of TBLV-induced tumors in B6 mice is dependent on MMTV Rem expression, but that proviral mutagenesis is dependent on mouse strain.

## RESULTS

### Accelerated T-cell tumors induced by lymphomagenic MMTV (TBLV) in the absence of Rem and mature B cells

Our previous results using splice donor site mutants of MMTV and TBLV in BALB/c mice indicated that MMTV-encoded Rem is involved in antagonizing proviral mutagenesis by Apobec family enzymes, including AID (18). Although Sag-mediated amplification of infected B and T lymphocytes is needed for efficient MMTV transmission and mammary cancer development (27, 30, 42), TBLV does not require Sag for the induction of T-cell lymphomas (38). Therefore, using Sag-independent TBLV, we tested the effect of Rem loss in multiple strains of genetically engineered mice that are available on the B6 background. We inoculated TBLV-WT and TBLV-SD independently into B6 mice. As observed for TBLV-induced tumors in BALB/c mice (18), no statistically significant difference in tumor incidence or latency was observed in wild-type B6 mice inoculated with TBLV-WT or Rem-null TBLV-SD (Fig. 2A). However, most TBLV-WT-inoculated mice developed T-cell lymphomas whereas, at the same dose, we observed a tumor incidence of 30-50% in BALB/c mice (18, 38). All TBLV-SD-injected animals developed tumors, but there was a trend toward acceleration of T-cell tumors in the absence compared to the presence of Rem (p = 0.076).

**Fig. 2.**
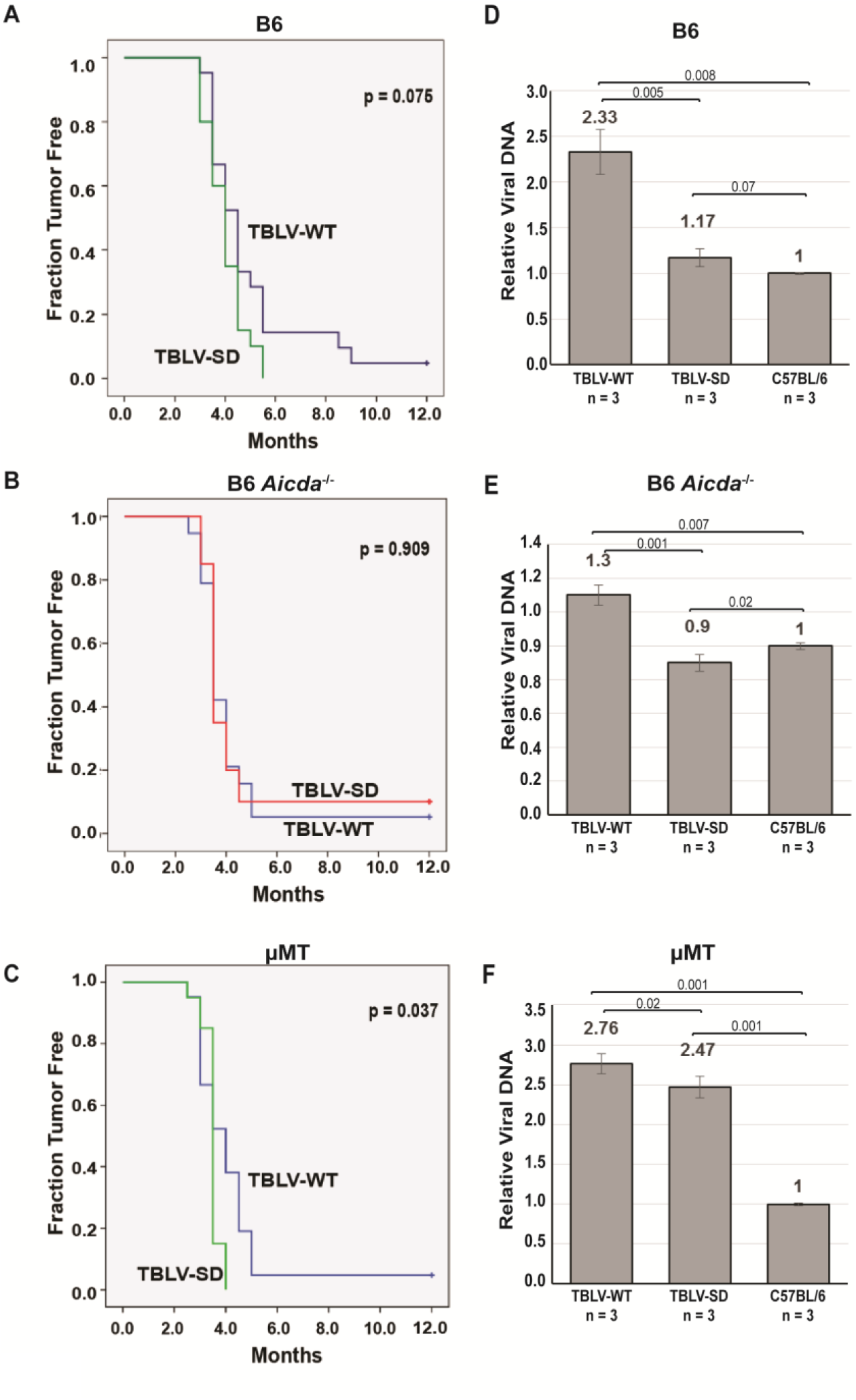
T-cell tumor development after high-dose infection reveals that TBLV infection does not require mature B cells. (A-C) Wild-type B6, *Aicda^-/-^*, or µMT mice (panels A to C, respectively) were injected with TBLV-WT or TBLV-SD and followed for tumor development, including enlarged thymus, spleen, and lymph nodes. The results were analyzed by Kaplan-Meier plots. Only µMT mice showed a significant difference between T-cell lymphoma development by TBLV-WT and TBLV-SD at this viral inoculum (p<0.05) as determined by Mantel-Cox log-rank tests. TBLV-SD tumors were accelerated in µMT mice relative to the other two strains, but survival plots did not differ for TBLV-WT among the three strains. (D-F) Proviral loads were assessed in three tumors from each viral strain by semi-quantitative PCR, which allowed quantitation relative to the three diploid copies of endogenous *Mtv*s in B6 mice (panels D to F, respectively). Proviral load appeared to be lowest in *Aicda^-/-^* mice, yet proviral copies did not correlate with tumor latency. The relative proviral copy numbers appeared to be highest in µMT mice that lack mature B cells.

Since our previous data suggest that Rem acts as a Vif-like antagonist of the Apobec family member AID in BALB/c mice and leads to AID proteasomal degradation in tissue culture (18), we also infected *Aicda*^-/-^ mice on the B6 background with TBLV-WT or TBLV-SD. Like wild-type B6 mice, most animals developed T-cell lymphomas, and many had both spleen and lymph node involvement. However, no statistical difference in tumor incidence or latency was observed between mice inoculated with TBLV-WT relative to those inoculated with TBLV-SD (p = .909) (Fig. 2B). Therefore, at this virus inoculum, the absence of AID eliminated the modest effect of Rem on TBLV-induced T-cell tumors in B6 mice.

To determine if TBLV, like MMTV, requires replication in mature B cells, B6 µMT (immunoglobulin heavy chain-knockout) mice and wild-type B6 mice were infected with TBLV-WT or TBLV-SD. The µMT strain was developed by introducing a neomycin-resistance cassette to disrupt one of the membrane exons of the IgM heavy chain gene (43). No expression of membrane-bound IgM is detectable and peripheral B-2 cells are lacking in this strain, although innate B-1 cells are made and produce IgE and IgG (44). TBLV-WT induced tumors in µMT mice with ∼100% efficiency (Fig. 2C), and the latency was not significantly different from wild-type B6 mice (Fig. 2A). Our results indicated that TBLV, a Sag-independent MMTV strain, does not require mature B cells for T-cell lymphomagenesis. We also assessed whether inoculation of TBLV-SD (lacking Rem expression) into µMT B6 mice would affect tumor incidence or latency compared to TBLV-WT. TBLV-SD infection led to a statistically accelerated appearance of T-cell lymphomas compared to those induced after TBLV-WT infection (p= 0.037) (Fig. 2C). These data suggested that the loss of Rem expression by TBLV provided a selective advantage for virus replication and/or tumorigenesis in µMT mice.

### Decreased TBLV proviral loads in the absence of Rem dependent on the presence of mature B cells

To determine if Rem expression affected proviral DNA levels, we extracted DNA from three independent TBLV-WT-induced tumors as well as the same number of TBLV-SD-induced tumors for analysis by PCR. Because B6 mice contain three endogenous *Mtv* proviruses with extensive sequence similarity to TBLV (45, 46), semi-quantitative PCRs that generated larger products allowed us to quantitate levels of haploid exogenous virus integrations relative to the diploid endogenous proviruses within tumor DNA (18). TBLV-SD-induced tumors in B6 mice had a proviral load that was statistically lower compared to tumors induced by TBLV-WT (Fig. 2D). These results suggested that the absence of Rem restricted TBLV replication prior to or during replication in T cells. Interestingly, proviral loads in *Aicda*^-/-^ mice were lower than in wild-type B6 mice in TBLV-WT-induced tumors (Fig. 2E), consistent with decreased replication of Rem-expressing virus in this strain. In contrast, proviral loads in TBLV-WT-induced tumors from µMT mice were similar to those induced in B6 mice (compare Figs. 2D and 2F), and the proviral load was twice that observed in *Aicda*^-/-^ mice (Fig. 2E). Interestingly, the TBLV-SD proviral load in µMT tumors was ∼2-fold increased over that observed in either wild-type or *Aicda*^-/-^ mice. These data are consistent with the idea that the absence of mature B cells increases TBLV replication.

### TBLV mutational profiles in the absence of Rem

Apobec family members are known restriction factors for retroviruses, including MMTV (6, 18). Proviral loads and transition mutations in WR<u>C</u> and TY<u>C</u> motifs typical of AID and mA3, respectively, were increased in BALB/c mammary tumors induced by MMTV lacking Rem expression. This difference was eliminated by induction of mammary tumors in AID-deficient *Aicda*^-/-^ mice, suggesting that Rem protects against multiple Apobec cytidine deaminases (18). In addition, previous high throughput sequencing of proviral DNA from BALB/cJ tumors showed a large increase in transition mutations in proviruses from TBLV-SD-induced tumors compared to those induced by TBLV-WT (18). The frequency of C-to-T mutations on the proviral plus strand (average number of mutations/clone) showed a dramatic increase from 0.08 to 0.73 (∼9-fold) when comparing TBLV-WT to TBLV-SD proviruses lacking Rem expression isolated from BALB/c tumors (18).

To assess whether proviral load differences between T-cell tumors induced by TBLV-WT and TBLV-SD were associated with alterations in Apobec-mediated hypermutation, tumor DNAs were amplified using TBLV-specific primers spanning the *env*-LTR region. Individual PCR products were cloned and analyzed by Sanger sequencing. In contrast to TBLV-induced BALB/c tumors (18), the average frequency of C-to-T mutations was similar for TBLV-WT and TBLV-SD proviruses (0.72 and 1.04 mutations/clone, respectively) obtained from B6 tumors (Table 1). These results suggested that MMTV-encoded Rem has little effect on Apobec-mediated proviral mutations in tumors induced in B6 relative to BALB/c mice.

**Table 1.**
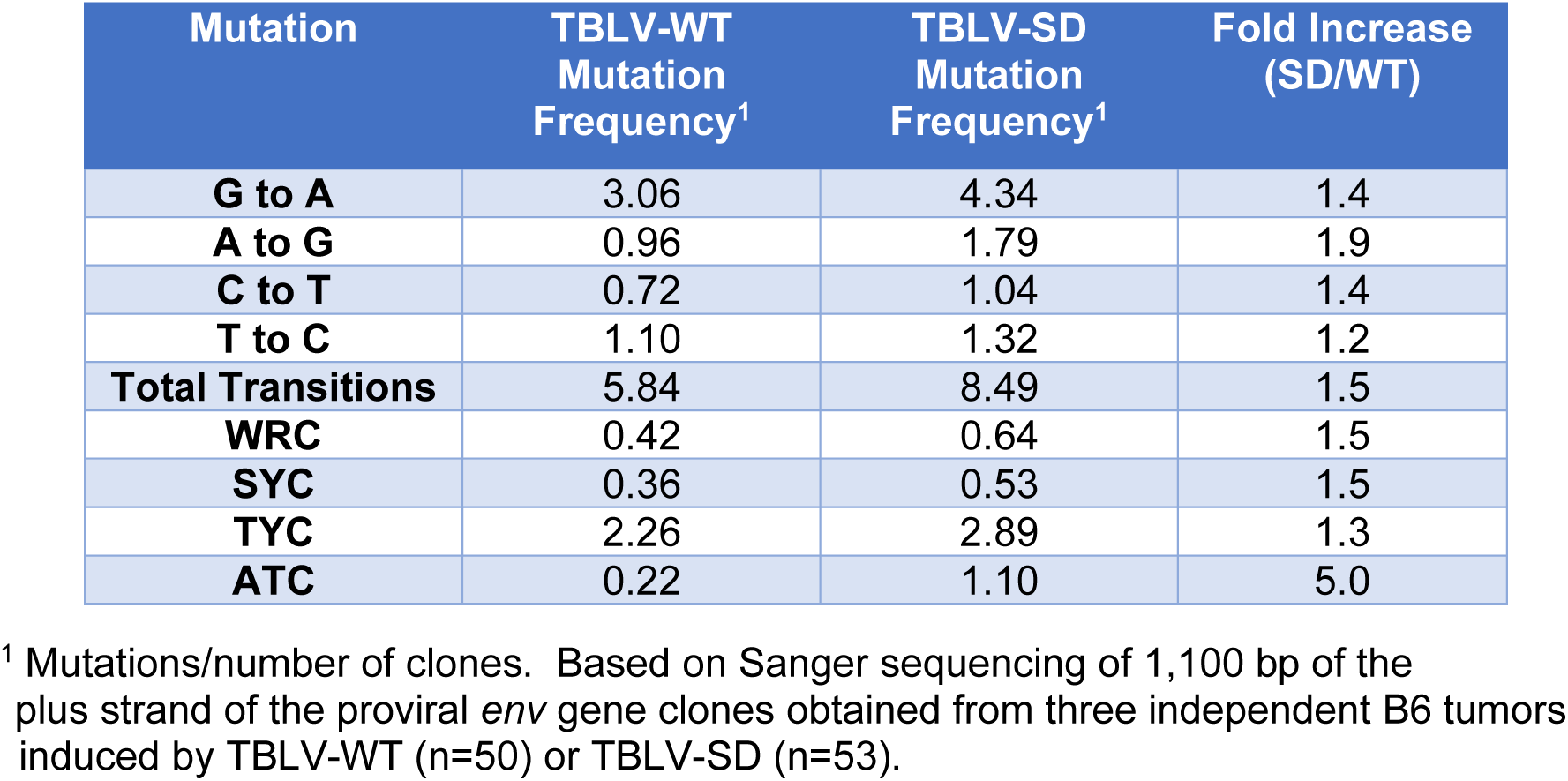
Mutation frequency in TBLV-WT and TBLV-SD proviruses from B6 T-cell tumors by Sanger sequencing.

Since the sequence context of Apobec-induced proviral mutations often reveals the identity of the enzyme involved (47), we analyzed mutations within specific sequence motifs. Analysis of mutations typical of murine AID (WR<u>C</u> motif) indicated a small increase for the TBLV-SD envelope region relative to TBLV-WT proviruses (1.5-fold) in B6 tumors (Table 1). In contrast, WR<u>C</u>-motif (W=A or T, R=A or G) mutations were increased by 2.6-fold in TBLV-SD compared to TBLV-WT proviruses obtained from BALB/c tumors (18).

Proviral mutations typical of mA3 (TY<u>C</u>; Y=T or C) were only 1.3-fold higher in TBLV-WT compared to TBLV-SD proviruses isolated from B6 tumors (Table 1), whereas mutations were 2-fold higher in TBLV-SD versus TBLV-WT proviruses from BALB/c tumors (18). SY<u>C</u>-motif (S=C or G, Y = T or C) mutations, which have been associated with AID enzymatic activity at lower frequency on immunoglobulin genes (48, 49), were 4.9-fold higher for TBLV-SD relative to TBLV-WT proviruses in BALB/c mice (18). However, this value was only 1.5-fold greater in TBLV-SD proviruses compared to TBLV-WT proviruses in B6 mice (Table 1). The average frequency of AT<u>C</u> motif mutations, which have been associated with mA3 activity *in vitro* (50), was increased 2.6-fold for TBLV-SD relative to TBLV-WT proviruses in BALB/c mice (0.47 versus 0.18 mutation/clone, respectively) (18). A 5-fold increase of AT<u>C</u>-motif mutations was observed in proviruses isolated from B6 mice (1.10 versus 0.22 mutations/clone for TBLV-SD and TBLV-WT, respectively) (Table 1). Therefore, the proviral mutations most affected by the absence of Rem in B6 tumors were those associated with the AT<u>C</u> motif, although the frequency of cytidine mutations remained highest in the TY<u>C</u> motif for either TBLV-WT or TBLV-SD. These data suggested that mutations in the AT<u>C</u> and TY<u>C</u> motifs are not produced by the same enzymes. Furthermore, the difference in proviral mutations observed between TBLV-WT and SD-induced BALB/c and B6 tumors was dramatically different.

### Apobec-mediated mutations in TBLV proviruses depend on mouse genetic background

The distribution of cytidine mutations was compared by Sanger sequencing of individual proviral clones from tumors induced in wild-type B6 as well as *Aicda*^-/-^ mice. G-to-A mutations were not increased in proviruses from *Aicda*-insufficient mice infected with TBLV-SD relative to those in TBLV-SD-infected mice (Table 2). Scatter plot comparisons between tumors from wild-type mice revealed that only AT<u>C</u>-motif proviral mutations were increased after infection with TBLV lacking Rem expression. However, TY<u>C</u>-motif mutations were the most abundant in proviruses from both TBLV-WT and SD-infected mice (Fig. 3A). Analysis of the distribution of AT<u>C</u>-motif mutations also revealed a significant increase within proviruses from tumors obtained from *Aicda*^-/-^ mice infected with TBLV-SD (Rem-null) relative to TBLV-WT (Fig. 3B). Surprisingly, a significant increase was observed in WR<u>C</u>-motif mutations between TBLV-SD and TBLV-WT proviruses obtained from tumors lacking AID expression. These results suggest that AID is not the sole enzyme responsible for proviral mutations in WR<u>C</u> motifs.

**Fig. 3.**
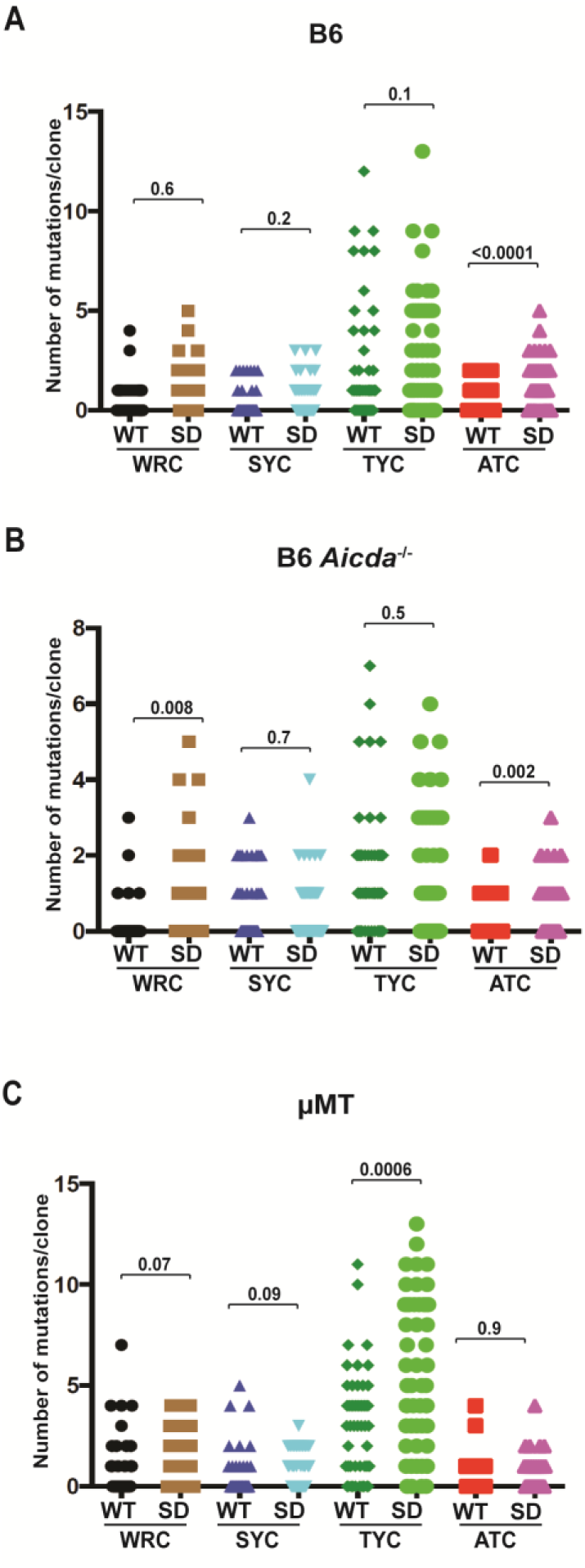
Apobec-associated proviral mutations in tumors induced by high doses of TBLV-WT and TBLV-SD. Independent cloned sequences were obtained from three tumors in different animals. The number of cytidine mutations within different motifs on either proviral strand is given for each clone. WR<u>C</u> and SY<u>C</u>-motif mutations have been associated with AID expression, whereas TY<u>C</u> and AT<u>C</u>-motif mutations have been linked to mA3 expression. Statistical significance by non-parametric Mann-Whitney tests is indicated on the scatter plots. (A) Comparison of the distribution of mutations within the proviral envelope gene from TBLV-WT or TBLV-SD-induced tumors from wild-type B6 mice. (B) Comparison of the distribution of mutations within the proviral envelope gene in TBLV-WT or TBLV-SD-induced tumors from *Aicda*^-/-^ mice. (C) Comparison of the distribution of mutations within the proviral envelope gene from TBLV-WT or TBLV-SD-induced tumors from µMT mice lacking mature B cells.

**Table 2.** Mutation frequency in TBLV-WT and TBLV-SD proviruses from B6 *Aicda*^-/-^ T-cell tumors by Sanger sequencing.

| Mutation | TBLV-WT<br>Mutation<br>Frequency<sup>1</sup> | TBLV-SD<br>Mutation<br>Frequency<sup>1</sup> | Fold Increase<br>(SD/WT) |
| --- | --- | --- | --- |
| **G to A** | 2.38 | 2.09 | 0.9 |
| **A to G** | 0.58 | 0.60 | 1.0 |
| **C to T** | 0.22 | 0.35 | 1.6 |
| **T to C** | 0.78 | 0.65 | 0.8 |
| **Total Transitions** | 3.96 | 3.69 | 1.0 |
| **WRC** | 0.16 | 0.60 | 3.8 |
| **SYC** | 0.54 | 0.56 | 1.0 |
| **TYC** | 1.58 | 1.42 | 0.9 |
| **ATC** | 0.16 | 0.55 | 3.4 |

<sup>1</sup> Mutations/number of clones. Based on Sanger sequencing of 1,100 bp of the plus strand of the proviral *env* gene clones obtained from three independent B6 AID-KO tumors induced by TBLV-WT (n=50) or TBLV-SD (n=55).

The distribution in specific cytidine mutations also was examined in TBLV proviruses obtained from µMT tumors. The G-to-A mutations on the TBLV-SD plus strand were 2-fold higher than those on the plus strand of TBLV-WT proviruses (Table 3). Unlike proviruses from wild-type B6 and *Aicda*^-/-^ mice, AT<u>C</u>-motif proviral mutations were similar in tumors induced by TBLV-WT and TBLV-SD (Fig. 3C). Furthermore, TY<u>C</u>-motif proviral mutations were significantly increased in TBLV-SD-induced tumors relative to those induced by TBLV-WT. Since the distributions of proviral TY<u>C</u> and AT<u>C</u>-motif mutations are dissimilar between tumors induced by TBLV-SD and TBLV-WT in all three backgrounds examined, our data are consistent with the idea that different enzymes are responsible. Furthermore, the increase in TY<u>C</u>-motif mutations associated with mA3 activity (51) only in µMT tumor-derived proviruses suggests that these mutations occur after infection of immature B cells.

**Table 3.** Mutation frequency in TBLV-WT and TBLV-SD proviruses from B6 µMT T-cell tumors by Sanger sequencing.

| Mutation | TBLV-WT<br>Mutation<br>Frequency<sup>1</sup> | TBLV-SD<br>Mutation<br>Frequency<sup>1</sup> | Fold Increase<br>(SD/WT) |
| --- | --- | --- | --- |
| **G to A** | 3.74 | 7.59 | 2.0 |
| **A to G** | 0.74 | 0.57 | 0.8 |
| **C to T** | 0.54 | 0.43 | 0.8 |
| **T to C** | 0.98 | 0.45 | 0.5 |
| **Total Transitions** | 6.00 | 9.04 | 1.5 |
| **WRC** | 0.72 | 1.02 | 1.4 |
| **SYC** | 0.52 | 0.69 | 1.3 |
| **TYC** | 3.10 | 5.61 | 1.8 |
| **ATC** | 0.40 | 0.41 | 1.0 |

<sup>1</sup> Mutations/number of clones. Based on Sanger sequencing of 1,100 bp of the plus strand of the proviral *env* gene clones obtained from three independent B6 μMT tumors induced by TBLV-WT (n=50) or TBLV-SD (n=51).

### Lack of Rem expression accelerates wild-type B6 and AID-insufficient tumors after low-dose infection

To determine whether differences between tumor susceptibility of B6 mice to TBLV-WT and TBLV-SD were dose-dependent, we injected a 20-fold lower dose into both wild-type B6 and *Aicd*a^-/-^ mice. At this dose, we achieved approximately the same tumor incidence in BALB/c and B6 mice injected with TBLV-WT. The results were plotted by the Kaplan-Meier method (Fig. 4A). In contrast to the higher dose, TBLV-SD-induced tumors were accelerated compared to those appearing in TBLV-WT-injected mice in both mouse strains (p<0.001). No statistical difference in tumor latency or incidence was observed in comparisons between TBLV-WT or TBLV-SD in the two strains. These data argue that Rem interferes with TBLV-induced tumor development independent of AID expression.

**Fig. 4.**
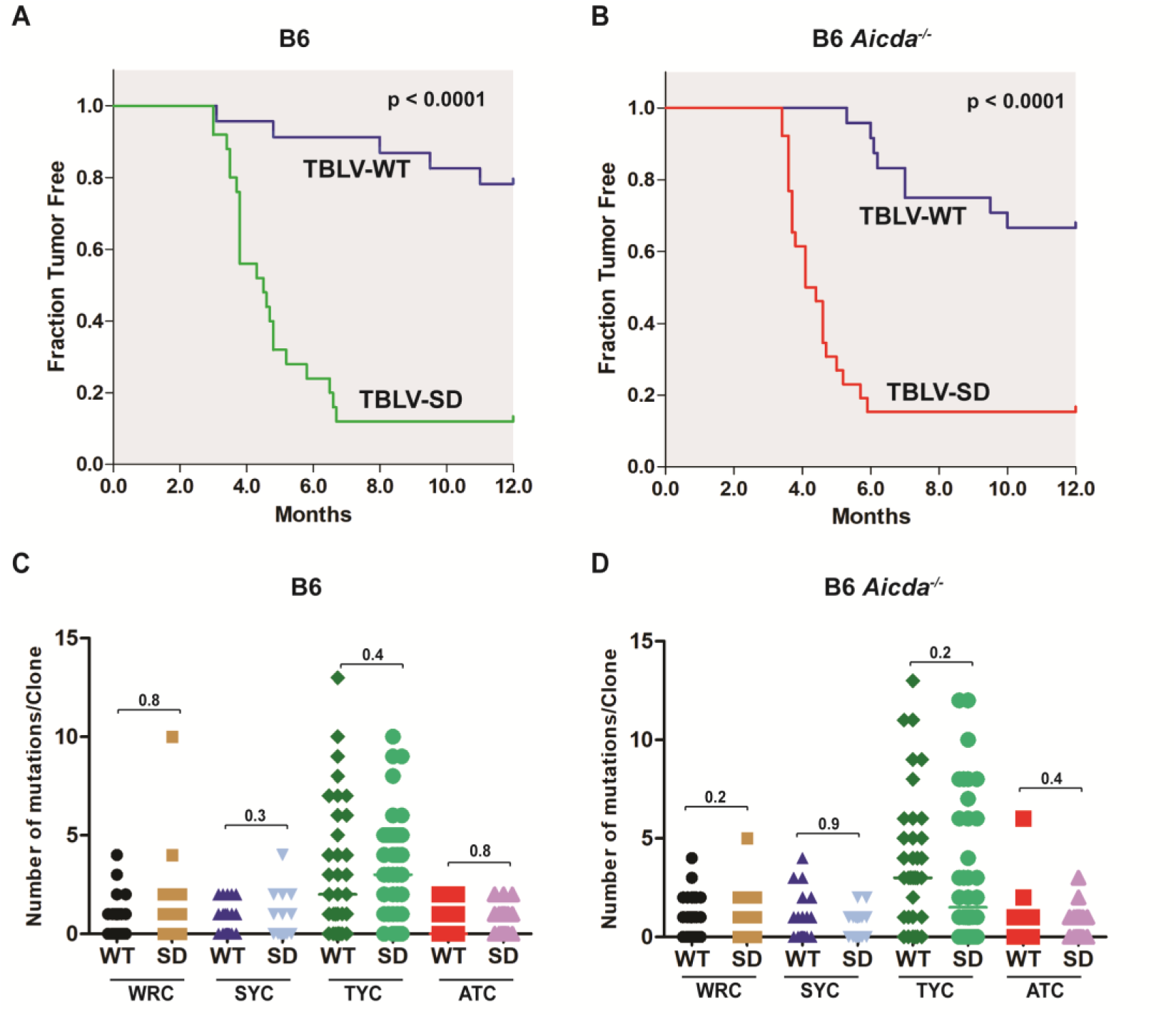
Low-dose infections in either wild-type or *Aicda^-/-^*B6 mice reveal accelerated tumorigenesis by TBLV in the absence of Rem expression. (A) Kaplan-Meier plots of tumors induced by the same low dose of TBLV-WT or TBLV-SD in wild-type mice on the B6 background. This dose was 20-fold lower than that used for results shown in Fig. 2. (B) Kaplan-Meier plots of tumors induced by the same low dose of TBLV-WT or TBLV-SD in *Aicda^-/-^* B6 mice. The p-values were calculated by Mantel-Cox log-rank tests. (C) Sanger sequencing of individual TBLV proviral clones from low-dose tumors in wild-type B6 mice. Differences in the distribution of cytidine mutations within the envelope gene of cloned proviruses (∼25 clones from three different tumors) were compared for WR<u>C</u>, SY<u>C</u>, TY<u>C</u>, and AT<u>C</u> motifs using scatter plots. The p-values were calculated by non-parametric Mann-Whitney tests. The most prevalent mutations were in the TY<u>C</u> motif typical of mA3. (D) Sanger sequencing of individual TBLV proviral clones from low-dose tumors in *Aicda^-/-^*B6 mice. Comparisons were performed as described in panel C.

To determine whether the difference in tumor latency between TBLV-WT and SD was due to Apobec-mediated mutagenesis, clones from the proviral region spanning the *env*-LTR region were obtained from three different B6 and *Aicda*^-/-^ tumors and subjected to Sanger sequencing. Independent clones then were analyzed for cytidine mutations in specific sequence contexts within the *env* region. WR<u>C</u>-motif mutations were detectable at low levels, but their distributions were not different between either TBLV-WT or SD proviruses within or between wild-type and knockout mice (Figs. 4C and 4D). In addition, the numbers of WR<u>C</u>-motif mutations/clone were similar in the presence or absence of a functional *Aicda* gene. Therefore, the cytidine mutations in WR<u>C</u> motifs are likely due to a different enzyme.

As noted in high-dose tumors, tumors derived from lower-dose infections had the most abundant proviral mutations in the TY<u>C</u> context typical of mA3. Similar to cytidine mutations in the WR<u>C</u> context, no statistical differences in the distribution of TY<u>C</u>-motif mutations were detected between wild-type and Rem-null proviruses in either B6 or *Aicda*^-/-^ mice (Figs. 4C and 4D). Also, unlike our results with MMTV-induced tumors in BALB/c mice (18), the numbers of TY<u>C</u>-motif mutations/clone did not decline in the absence of AID expression. No significant differences were observed in the SY<u>C</u> or AT<u>C</u> contexts in the presence or absence of Rem or AID expression. These results suggest that Apobec-mediated hypermutations are not responsible for tumor latency differences observed between TBLV-WT and TBLV-SD infections and are mouse-strain-dependent.

To increase the number of proviruses analyzed for Apobec-mediated changes, we performed MiSeq sequencing on the PCR products spanning the *env*-LTR region of proviruses derived from tumors appearing after the high-dose TBLV infection (Fig. 5A). As expected, analysis revealed that G-to-A mutations typical of Apobecs were the most abundant changes, but no significant differences were observed between TBLV-WT and SD proviruses in either wild-type or *Aicda*-insufficient B6 mice. We also analyzed tumors from high-dose infections by Nanopore sequencing with similar results (Fig. 5B). These results further confirm that the absence of Rem had no demonstrable effect on TBLV proviral mutations in tumors induced in B6 mice.

**Fig. 5.**
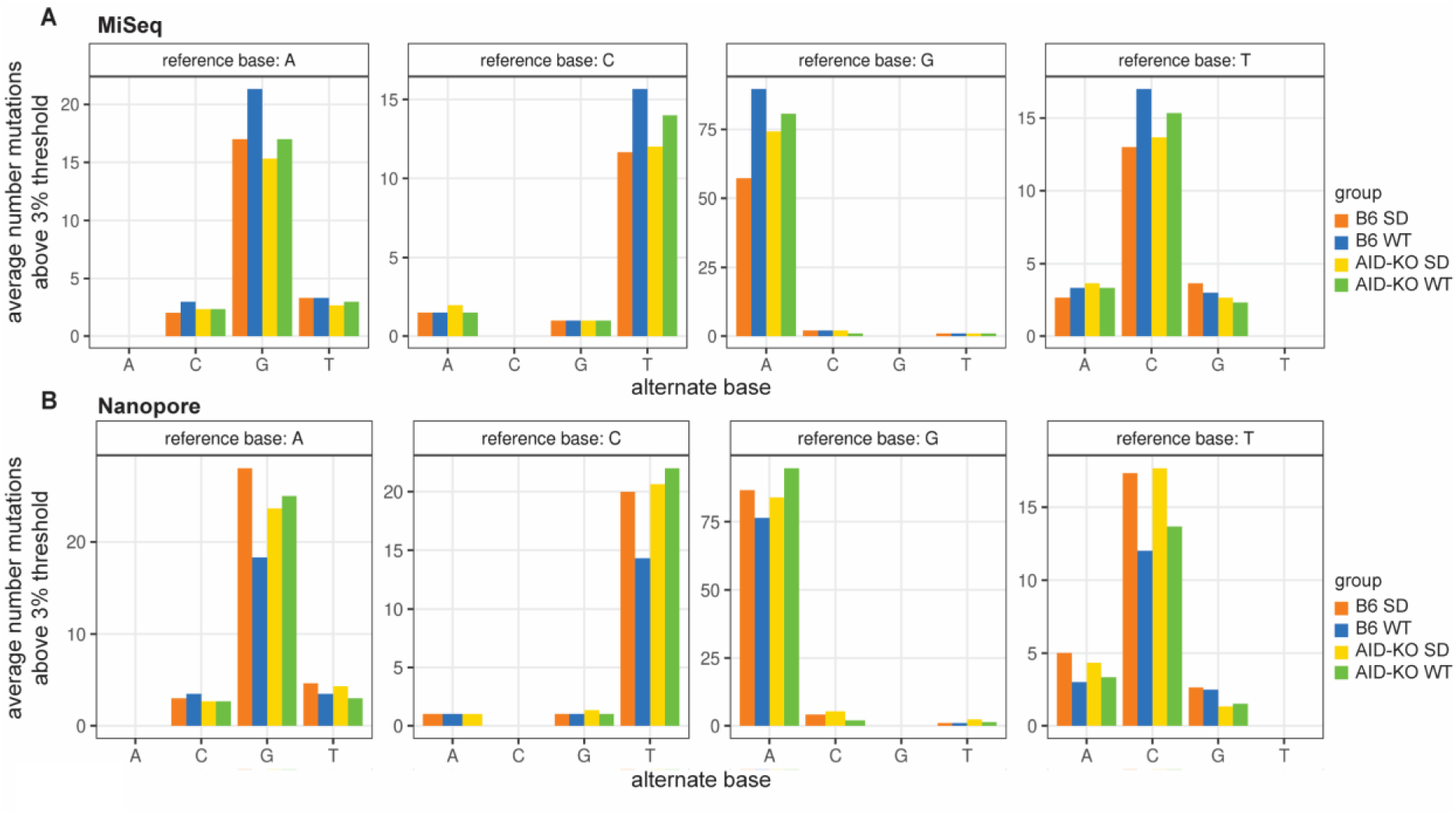
High-throughput sequencing of proviruses from tumors induced by low-dose infections of TBLV-WT and TBLV-SD. (A) MiSeq sequencing of the *env*-LTR proviral junction fragments obtained by PCR using DNA from TBLV-SD or TBLV-WT-induced thymic lymphomas of infected wild-type or AID-knockout mice. Tumors were obtained after high-dose injections. Each bar represents results from three independent tumors. The average number of mutations above a 3% threshold are plotted for each position relative to the cloned TBLV sequence. Pairwise comparisons between groups showed no statistical differences. (B) Nanopore sequencing of the *env*-LTR proviral junction fragments obtained by PCR using DNA from TBLV-SD or TBLV-WT-induced thymic lymphomas of infected wild-type or AID-knockout mice. Tumors were obtained after low-dose injections.

### Analysis of LTR enhancer repeats within tumor-derived TBLV proviruses

Acceleration of T-cell lymphomas was observed after infection with TBLV-SD compared to TBLV-WT infections in either wild-type or *Aicda*^-/-^ B6 mice. We considered whether differential selection for *cis*-acting LTR elements, which are known to affect TBLV disease specificity (37, 52, 53), could accelerate TBLV-SD induction of tumors in µMT mice after Apobec-mediated mutagenesis. Specifically, TBLV proviruses have a triplicated T-cell enhancer element within the LTR that increases viral transcription in T cells (36, 52). Our previous work has shown that changes within this region are selected during tumor passage, with 1- and 4-enhancer elements being less active transcriptionally than 3-enhancer repeats (53). As expected, PCR performed with LTR-specific primers showed a predominance of 3 repeats within the LTR, consistent with the injected clonal virus (38). No change in the number of enhancer repeats within TBLV-SD proviruses relative to TBLV-WT proviruses was observed in tumors from wild-type or AID-deficient (*Aicda*^-/-^) B6 mice that would confer an obvious selective advantage (Figs. 6A and B). However, the results revealed that the viral populations within tumors were mixed, consistent with LTR selection during lymphomagenesis (53).

**Fig. 6.**
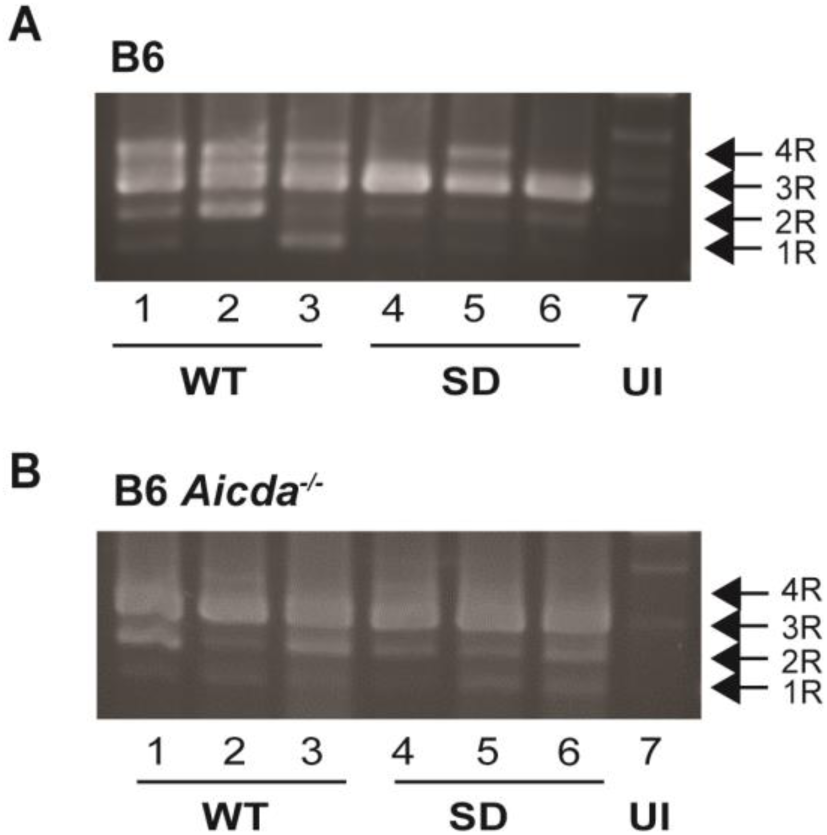
LTR enhancer repeats within TBLV proviruses obtained from low-dose tumors induced in wild-type and *Aicda^-/-^* mice. (A) Analysis of LTR enhancer repeats in tumors induced by TBLV-WT (WT) and TBLV-SD (SD) in wild-type B6 mice. PCRs were performed on three tumors from each viral strain prior to agarose gel electrophoresis. The positions of LTRs containing 1 (1R), 2 (2R), 3 (3R), or 4 (4R) copies of the 62-bp sequence in the TBLV enhancer (53) are indicated. The infectious TBLV-WT and TBLV-SD clones both have a 3R enhancer (38). (B) Analysis of LTR enhancer repeats in tumors induced by TBLV-WT and TBLV-SD in *Aicda^-/-^* mice on the B6 background. UI = DNA from uninfected mice. Faint background bands are observed in DNA from uninfected mice, presumably due to homology of the primers to the endogenous *Mtv*s.

### Differential mA3 and AID levels and signaling in B6 and BALB/c mice

To understand the correlation between AID and mA3 expression with proviral hypermutation in BALB/c and B6 tumors, we isolated splenocytes from at least three weanling mice of each strain. Pooled cells then were treated for up to 4 days with Concanavalin A (ConA) to stimulate T cells, with interleukin-4 (IL-4) and lipopolysaccharide (LPS) to stimulate AID production in B cells, and with interferon beta (IFNβ) to simulate a viral infection. After incubation with these stimulants, cell lysates were used for Western blotting to determine AID and mA3 levels (Fig. 7A). As expected, mA3 levels were higher in B6 splenocytes relative to those obtained from BALB/c mice, and only the Δexon5 isoform was expressed (50). Comparisons to unstimulated splenocyte extracts revealed that IL-4/LPS treatment gave a large increase in AID levels in both BALB/c and B6 cells, although slighter greater in BALB/c cells (compare lanes 4 and 8). In contrast, mA3 levels showed a modest increase in response to IL-4/LPS. Surprisingly, ConA-stimulated AID and mA3 levels in BALB/c cells were similar to those observed with IL-4/LPS (Fig. 7a, lanes 2 and 4). However, AID levels did not increase in B6 splenocytes in response to ConA (lane 6). Levels of mA3 showed a small increase similar to BALB/c splenocytes after ConA treatment (compare lanes 5 and 6 with lanes 1 and 2). IFNβ treatment gave no change in relative amounts of either mA3 or AID at this time point, but was active on BALB/c and B6 splenocyte populations as demonstrated by increases in phosphorylated Stat1 (Fig. 7B). Together, these results indicated that there are differences in signaling pathways for AID induction in BALB/c and B6 lymphocytes as well as differences in mA3 levels.

**Fig. 7.**
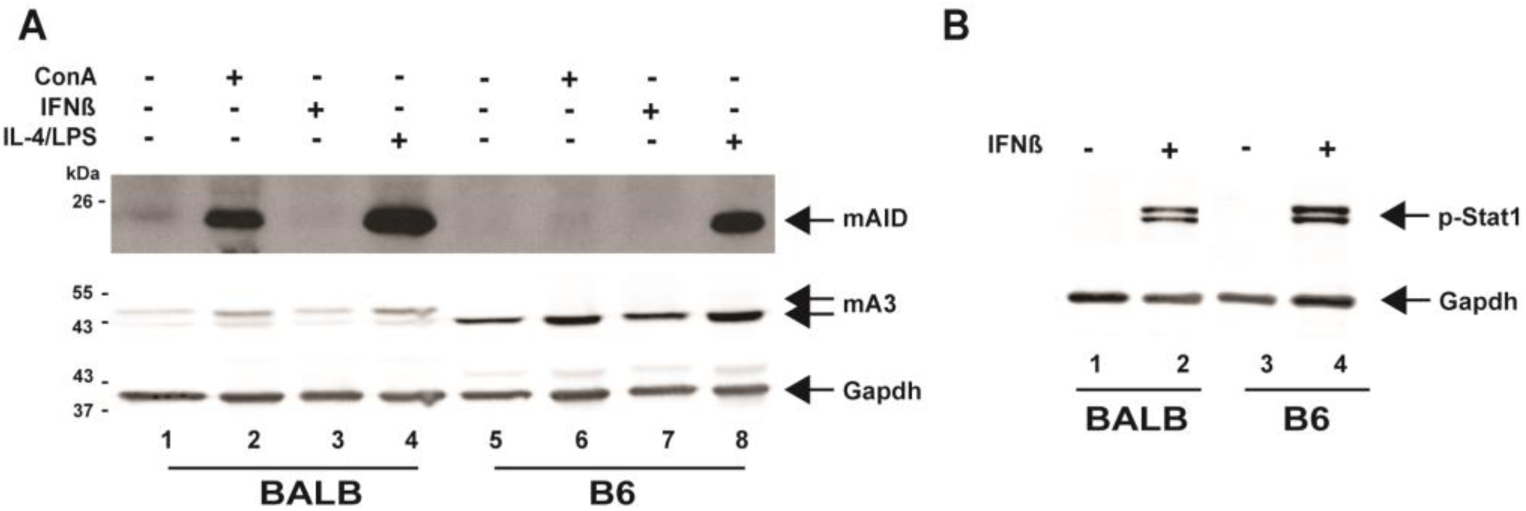
Expression of mAID and mA3 differs in B6 and BALB/c splenocytes after external ligand stimulation. (A) Splenocyte responses to Concanavalin A (ConA), IFNβ, or IL-4 plus LPS. Splenocytes from uninfected BALB/c or B6 mice were used for whole cell lysate preparation prior to culture (lanes 1 and 5), ConA (lanes 2 and 6), 200 U/ml IFNβ (lanes 3 and 7), or IL-4 plus LPS (lanes 4 and 8) for 4 days. Lysates were used for Western blotting and incubation with antibodies specific for mAID (top), mA3 (middle), or Gapdh (bottom). Note that BALB/c mice express two isoforms of mA3 mRNA (with and without exon 5), whereas B6 express predominantly the isoform without exon 5. (B) Treatment with IFNβ gives similar signaling responses in BALB/c and B6 splenocytes. Lysates from splenocytes treated with or without 1000 U/ml IFNβ for 5 h were used for Western blotting with antibodies specific for phosphorylated Stat1 (p-Stat1) or Gapdh. Two bands observed with p-Stat1 antibody correspond to phosphorylated Stat1α and Stat1β isoforms and are expressed in both BALB/c and B6 mice.

### Effects of Rem and AID on cytokine signaling

To determine differences in transcription that alter cell signaling induced in the presence and absence of Rem expression, we extracted RNA from three independent tumors induced by TBLV-WT as well as from TBLV-SD-infected mice. In tumors from infected wild-type B6 mice, >15,000 transcripts were detected by RNA-seq. Using a 2-fold difference in mRNA levels as a cutoff, 30 transcripts were significantly upregulated in TBLV-SD-induced tumors relative to those induced by TBLV-WT. Nine transcripts, including *Ighg3* and *C4b,* were significantly downregulated (Fig. 8A). Analysis of tumors from *Aicda*^-/-^ mice infected with either TBLV-WT or TBLV-SD revealed a similar number of total mRNAs as those obtained from infected B6 tumors. In contrast to results with wild-type B6 mice, 424 different gene transcripts were significantly increased by at least 2-fold in AID-insufficient tumors infected with TBLV-SD relative to those infected with TBLV-WT (Fig. 8B). Three of the most highly upregulated transcripts in the absence of Rem expression were *Il2ra*, *Socs3*, and *Stat4*. In addition, 83 mRNAs were significantly downregulated in TBLV-SD-induced tumors when compared to mRNAs in TBLV-WT-induced tumors. The complete list of transcripts and their relative abundance after TBLV infection in B6 and *Aicda*^-/-^ mice is provided in Tables S1 and S2, respectively. Therefore, the most dramatic effect on the transcriptome of TBLV-induced tumors occurred in the absence of both AID and Rem expression.

**Fig. 8.**
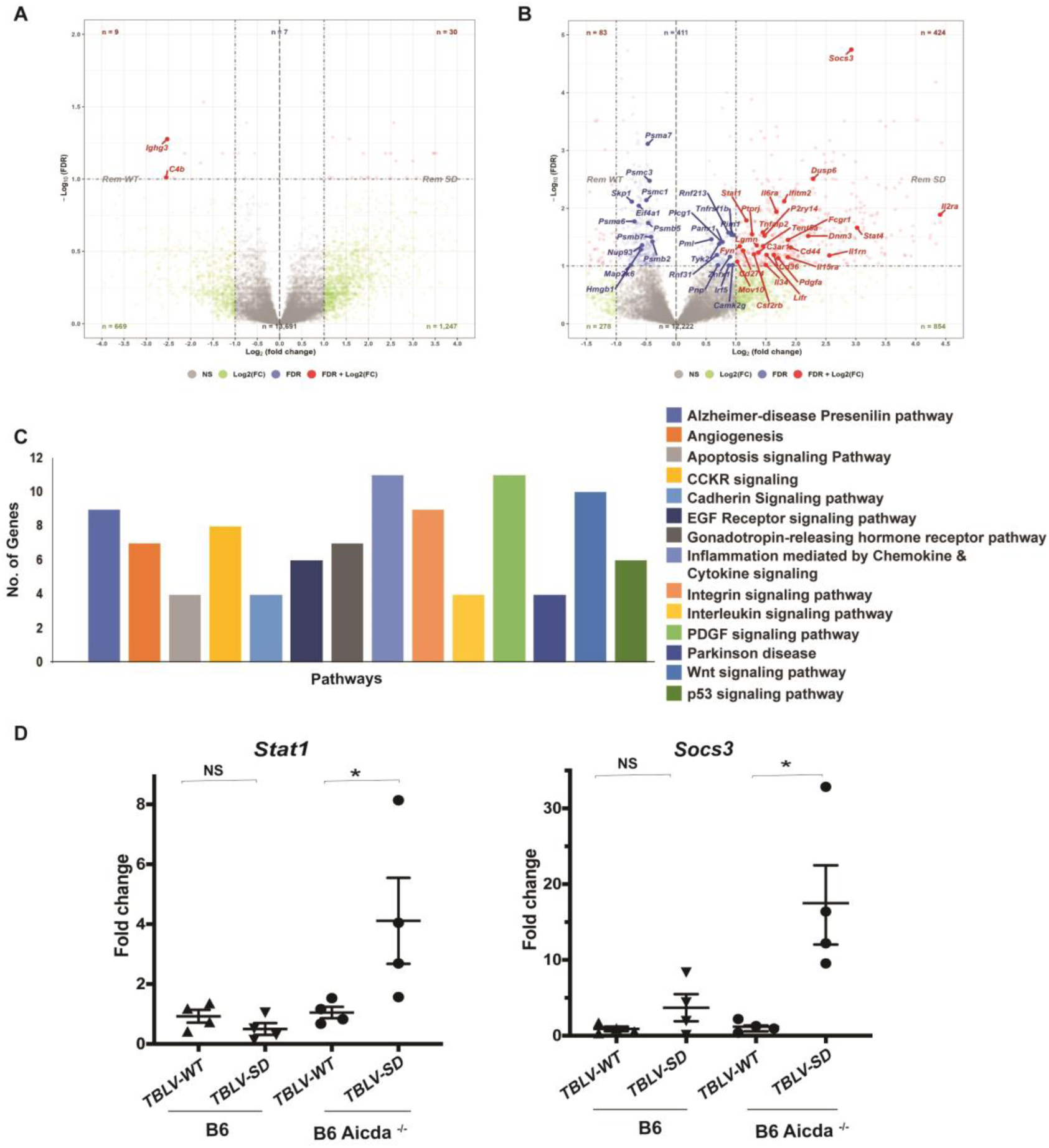
RNA-seq analysis indicates increased immune-related transcripts in the absence of Rem and AID expression. (A) Volcano plot comparing different transcripts from TBLV-WT and TBLV-SD-infected tumors from B6 mice. The false discovery rate (FDR) (-log_10_) was plotted versus the log_2_-fold change (FC) in mRNA abundance. Blue dots indicate transcripts that were significantly different in TBLV-SD-induced tumors. Red dots indicate transcripts that were significantly different in TBLV-SD-induced tumors and were changed more than 2-fold. Non-significant differences are shown by gray dots. The identities of some transcripts are provided. (B) Volcano plot comparing different transcripts from TBLV-WT and TBLV-SD-infected tumors from B6 *Aicda^-/-^* mice. (C) PANTHER analysis of differentially expressed genes in tumors induced by TBLV-SD relative to TBLV-WT. (D) Validation of *Stat1* and *Socs3* mRNAs expressed in wild-type and AID-insufficient B6 mice in tumors induced by TBLV-WT and TBLV-SD (4 tumors each). Each symbol represents RNA from a single tumor. Mean and range of values obtained by RT-qPCR are shown. NS = non-significant

To determine whether the genes upregulated in the absence of Rem and AID were associated with a particular cellular pathway, we used PANTHER analysis (www.pantherdb.org). The results revealed that >20 upregulated genes were associated with platelet-derived growth factor (PDGF) and chemokine/cytokine signaling (Fig. 8C). Although Rem functions both in Apobec antagonism and control of viral mRNA nuclear export and expression (18–20), these activities targeted post-transcriptional activities, not mRNA levels.

To validate the results of the RNA-seq analysis, we selected two genes, *Socs3* and *Stat1*, in the *Aicda*^-/-^ tumors induced by TBLV-SD (Rem-null) for further testing. Four tumors each from TBLV-WT and TBLV-SD-infected B6 and *Aicda*^-/-^ mice were used for reverse-transcription real-time PCR (Fig. 8D). As expected, both *Socs3* and *Stat1* mRNAs were increased relative to glyceraldehyde-3-phosphate dehydrogenase (*Gapdh*) levels in TBLV-SD-infected *Aicda*-deficient tumors relative to those in tumors from TBLV-WT-infected animals. However, neither *Socs3* nor *Stat1* mRNAs were statistically elevated in tumors induced by TBLV-SD relative to those induced by TBLV-WT in wild-type B6 mice. These results suggest an interplay between Rem and AID activities that affects RNA levels of multiple genes, including those involving cytokine signaling.

## DISCUSSION

We previously showed that infectious MMTV and TBLV proviruses lacking Rem expression obtained from tumors on the BALB/c background have multiple transition mutations typical of Apobec cytidine deaminases (18). The mutational differences between Rem-expressing and Rem-null proviruses were primarily in cytidine-containing motifs typical of AID and mA3 and were abolished in *Aicda*^-/-^ BALB/c mice (18). However, most single or double-knockout Apobec strains are on a B6 background, and previous experiments have shown that MMTV has higher viral loads in B6 *mA*3^-/-^ compared to wild-type mice (6). Therefore, we conducted experiments in B6 and Apobec-mutant mice to confirm the mutational phenotype observed in BALB/c mice. Surprisingly, only small mutational differences were observed between TBLV-SD (Rem-null) and TBLV-WT proviruses from B6 tumors.

Because B6 mice are unable to efficiently present C3H MMTV Sag due to a mutation in an MHC class II gene (54), the Sag-independent MMTV strain, TBLV (36, 38), was used for infection of B6 mice. BALB/c and B6 mouse strains are immunologically distinct (55). In response to pathogens, T cells from B6 mice preferentially produce Th1 cytokines with high interferon-gamma (IFNγ) and low interleukin (IL-4), whereas those from BALB/cJ produce Th2 cytokines with low IFNγ and high IL-4 production (35). BALB/c mice also have only a transient antibody response to MMTV infection (56). Therefore, we anticipated that B6 mice would be more resistant to TBLV-induced tumors. Surprisingly, the TBLV-WT dose that gave a 30-50% tumor incidence in BALB/cJ mice (18, 38) resulted in a nearly 100% incidence in B6 mice (Fig. 1). In contrast, a 20-fold lower TBLV-WT dose produced approximately the same tumor incidence in BALB/c and B6 mice (Fig. 5).

We also analyzed TBLV-induced tumors in B6 mice lacking mature B cells (µMT) (43) or AID expression (*Aicda*^-/-^) (41). At the higher dose, TBLV-SD (Rem-null)-infected µMT mice had a significantly shorter tumor latency compared to TBLV-WT-infected animals, whereas AID-knockout mice showed no significant difference. However, at the lower dose, both AID-knockout and wild-type B6 mice had increased tumor incidence and latency after infection with TBLV-SD relative to TBLV-WT. These results suggest that Rem expression inhibits tumor formation, which often reflects proviral load since TBLV and MMTV primarily induce tumors by multiple insertions. Only a few of these insertions activate specific proto-oncogenes that provide a cell growth advantage (57–60). Examination of proviral loads revealed that tumors in B6 mice had lower TBLV proviral loads in the absence of Rem expression, whereas no difference was observed in µMT mice. AID-knockout mice showed lower proviral loads between TBLV-WT and SD-induced tumors, yet overall loads were reduced compared to those of either B6 wild-type or µMT mice (Fig. 2). These results suggest that mature B cells, which are not required for TBLV transmission, inhibit virus replication. Decreased proviral loads in the absence of AID expression may result from effects on B-cell maturation and reduced targets for TBLV replication (61).

Surprisingly, we observed much smaller differences in the numbers of mutations/clone between TBLV-WT and TBLV-SD proviruses obtained from B6 tumors (Figs. 3-5) relative to those from BALB/c tumors (18). In BALB/c tumors, TBLV-SD proviruses (Rem-null) had increased TY<u>C</u> and WR<u>C</u>-motif mutations typical of mA3 and AID, respectively, relative to TBLV-WT proviruses (18). The numbers of mutations and differences between TBLV-WT and TBLV-SD proviruses largely were abolished in the absence of AID expression (18). However, increased mutations/clone and significantly different distributions only were observed in mA3-associated TY<u>C</u> motifs in TBLV-SD versus TBLV-WT proviruses in µMT mice, which lack mature B cells (43), and not in either wild-type B6 or *Aicda*-/- mice (Fig. 3). These data imply that mA3 expression is higher in immature B cells. In contrast, significant differences in the distribution of AT<u>C</u>-motif mutations, a minor proportion of the total, within TBLV-WT and SD proviruses only were observed in wild-type B6 and AID-knockout tumors. This difference in proviral mutations was not observed in µMT tumors. Although AT<u>C</u>-motif mutations have been attributed to mA3 from *in vitro* experiments (50), the differences in their distributions in the absence of Rem and in different mouse strains suggest that cytidine mutations within AT<u>C</u> and TY<u>C</u> sequences result from independent enzymes.

WR<u>C</u> and SY<u>C</u>-motif mutations both have been associated with AID-induced mutations of the immunoglobulin variable region genes, with mutations in WR<u>C</u> motifs being much more common (62). Although AID is known to be responsible for antibody affinity maturation in germinal center B cells, AID-catalyzed somatic hypermutation and class-switch recombination also have been demonstrated in immature B cells (61). We observed significant differences in the distribution of WR<u>C</u>-motif mutations between TBLV-WT and TBLV-SD-induced BALB/c tumors (18). This difference was not found in wild-type B6 tumors using Sanger sequencing of individual clones within the envelope region (Fig. 3). Unexpectedly, proviral mutations in WR<u>C</u> motifs were observed with a similar frequency and distribution after infection with TBLV-WT or Rem-null virus in wild-type B6 and µMT mice. However, in *Aicda*^-/-^ mice, WR<u>C</u>-motif mutations increased in the absence of Rem expression. SY<u>C</u>-motif mutations did not follow this pattern, consistent with the idea that different enzymes induced cytidine mutations in the WR<u>C</u> and SY<u>C</u> context (Fig. 3).

Replication of certain murine leukemia viruses (MuLVs) (gammaretroviruses) is inhibited by mA3 (63–65). B6 infections with a glyco-Gag mutant of Moloney MuLV (MoMuLV) reduced infectivity, but this was abolished in *mA3*^-/-^ mice (65). Hypermutations were not observed in the absence of glyco-Gag in infected wild-type B6 mice. Infection of BALB/c mice with a glyco-Gag defective mutant also showed decreased titers relative to wild-type mice, yet proviral hypermutations were not examined (65). Glyco-Gag is an N-terminal extension of the Gag structural capsid protein precursor that allows membrane insertion (66). MuLVs without glyco-Gag expression appear to have a late-stage defect that involves budding and release in cultured fibroblasts (67). Glyco-Gag is incorporated into virions to stabilize the capsid and prevents mA3 access to reverse transcripts (65). Inhibition of mA3 by glycosylated Gag does not require deaminase function (68). Nevertheless, certain MuLV strains, such as AKV, show hypermutation of the proviral genome even when glyco-Gag is expressed (63, 64, 69).

Co-transfection of mA3 from BALB/c (either exon5-plus or minus isoforms), NIH Swiss, or B6 mice into 293T cells together with AKV or MoMLV infectious clones revealed that AKV, not MoMuLV proviruses were hypermutated after infection of NIH3T3 cells. The AKV hypermutations primarily occurred in the TT<u>C</u> motif (63), which is consistent with our use of the TY<u>C</u> motif. AKV restriction correlated with mA3 levels rather than specific mRNA isoforms, with B6 neonatal splenocytes having the highest amounts of mA3 protein relative to BALB/c or NIH Swiss splenocytes. Both rat A3 and human A3G hypermutated MoMuLV. Purification of splenic B and T cells from B6 mice and infection with MoMuLV or AKV clones revealed mA3-induced mutations within both cell types, but only in AKV proviruses (63). These experiments suggested that the MuLV strain determined their susceptibility to mA3 hypermutation. In contrast, our results showed that the same viral strain (TBLV) displayed proviral mutations dependent on the mouse strain infected.

Low splenocyte levels of mA3 relative to AID correlated with tumor-associated proviral hypermutations in BALB/c mice, whereas the opposite was true in B6 mice (Fig. 7). Moreover, BALB/c and B6 splenocytes responded differently to some external signals. For example, ConA induced mA3 in both BALB/c and B6 splenocytes, whereas AID was induced only in BALB/c spleen cells (Fig. 7A). However, LPS binding to TLR4 leads to signaling through MyD88 (70) to yield increased AID levels in splenocytes from both mouse strains. Previous reports have shown that viruses, including MMTV, use LPS on bacterial cells to facilitate infection (71, 72), yet this signal likely will increase cytidine deaminase levels that curtail virus replication, necessitating Rem antagonism of Apobecs.

AID is known to alter gene expression epigenetically (73). Hematopoietic precursor/stem cells (HPSCs) lacking AID expression on the B6 background upregulate the expression of multiple genes, including those encoding the transcription factors Cepbα (74), Klf1 (75), and Gata1 (76). However, decreased gene expression is observed for suppressor of cytokine signaling 3 (Socs3) (77). *Aicda*-/- HPSCs showed skewing of hematopoiesis toward the myeloid lineage (73). Interestingly, our RNA-seq analysis revealed that *Socs3* mRNA was increased in the absence of both AID and Rem expression (Fig. 8), in addition to altered mRNA expression of IFN-induced genes and those involved in growth factor and cytokine signaling pathways. These data suggest that AID has effects on innate immunity and cytokine signaling that require further characterization and that Rem expression manipulates such pathways.

Why don’t proviral hypermutations correlate with TBLV-induced tumor latency? One explanation is that Apobec-mediated hypermutations reduce replication in T lymphocytes, but recombination with endogenous *Mtv*s allows selection for replication-competent TBLVs that cause insertional mutagenesis (18). Such viral recombinants have been observed in both MMTV-induced mammary tumors and TBLV-induced T-cell lymphomas (18, 78, 79). Another non-exclusive possibility is that the multiple gene products of Rem have separate effects on proviral mutation and tumor susceptibility. As suggested previously, uncleaved Rem is likely an AID antagonist (26), whereas SP is a Rev-like protein that facilitates MMTV mRNA export and expression (20, 24, 80). In contrast, the Rem C-terminal cleavage product, Rem-CT, has an unusual trafficking pathway through the ER and endosomes (26). SP is synthesized by both TBLV-WT and SD, suggesting that differences in gene expression between these viruses are due to uncleaved Rem and Rem-CT. Since uncleaved Rem co-expression likely leads to AID proteasomal degradation (18), we speculate that Rem-CT is responsible for suppressing tumor latency, perhaps through cytokine manipulation.

## MATERIALS AND METHODS

### Mouse infections

BALB/cJ and C57BL/6J mice were obtained from Jackson Laboratories (Bar Harbor, ME). *Aicda^-/^*^-^ mice on the B6 background were generated in the laboratory of Dr. Tasuku Honjo (41) and were kindly provided by Dr. Michel Nussenzweig (The Rockefeller University). For each strain of mice, age-matched weanlings 4-to-6 weeks of age were injected intraperitoneally with either 1 x 10^6^ (low dose) or 2 x 10^7^ (high dose) stably transfected Jurkat T cells. The infectious clones of TBLV-WT and TBLV-SD in a plasmid vector expressing hygromycin resistance have previously been described (38). Infectious clones were used for transfection of Jurkat cells and selected for three weeks in hygromycin and then screened for production of equivalent amounts of TBLV-WT or TBLV-SD (Rem-null) as judged by Western blotting with antibodies to MMTV capsid protein (p27) (56). Mice were monitored for the development of thymic and/or splenic tumors for a period of 9-12 months. All experimental procedures were approved by the Institutional Animal Care and Use Committee of The University of Texas at Austin.

### DNA isolation, cloning, and Sanger sequencing

Genomic DNA extracted from tumors induced by TBLV-WT or TBLV-SD was used for PCR with primers *env*7254(+) (5’-ATC GCC TTT AAG AAG GAC GCC TTC T-3’) and LTR9604(-) (5’-GGA AAC CAC TTG TCT CAC ATC-3’) for the region spanning the envelope-LTR junction. PCRs were performed in 25 µl with JumpStart RED Accutaq LA polymerase (Sigma-Aldrich, cat# D8045) in the supplied buffer, 500 ng of tumor DNA, 25 to 50 pmol of each primer, and 0.5 mM deoxynucleoside triphosphates. PCR parameters were: 94°C for 1 min, 10 cycles at 94°C for 10 sec, 53°C for 30 sec, 68°C for 2 min followed by 25 cycles of 95°C for 15 sec, 50°C for 30 sec, and 68°C for 2 min with a final incubation at 68°C for 7 min. Sequences encompassing the SD site and the *env* gene were acquired after cloning the PCR fragments inserted into pGEM-Teasy (Promega). Sanger sequencing with primers env7254(+) and env8506(-) (5’-GCA CTT GGT CAA GGC TCC TCG-3’) was used to verify clones as previously described (18). Sanger sequencing enabled identification of proviruses containing recombinants with endogenous *Mtv* proviruses.

### PCRs and RT-PCRs

To analyze the numbers of LTR-enhancer repeats, PCR was performed with 10 pmol of primers TBLV-LTR408(+) (5’- CCA ATA AGA CCA ATC CAA TAG GTA GAC -3’) and TBLV-LTR786(-) (5’- CAC TCA GAG CTC AGA TCA GAA C -3’), 100 or 200 ng of tumor DNA, and JumpStart^TM^ Taq ReadyMix (Sigma, cat# P0982). PCR parameters were: 94 °C for 5 min, then 35 cycles at 94 °C for 30 sec, 56 °C for 30 sec, 72 °C for 30 sec, and a final incubation at 72 °C for 7 min. Semi-quantitative PCR was performed with *Mtvr2* as the single copy gene standard using primers *Mtvr2*(+) (5’-TCT GGG ATC CGC TTC CTC AT-3’) and *Mtvr2*(-) (5’-CCA GTC CTT GGC CCT CAT TTA-3’). MMTV-specific primers *pol*4235(+) and *pol*5835(-) in the viral polymerase gene were used to measure proviral sequences (18).

The qRT-PCRs were performed in triplicate using a ViiA7 Real-time PCR System (Applied Biosystems). Briefly, 1 µg of total DNase I-treated RNA was reverse transcribed using the Superscript first-strand synthesis kit (Invitrogen). The cDNA was diluted in DNase-RNase free water, and 20 ng was used for the reaction. The expression of the *Gapdh* gene was used for normalization. The reaction mixture consisted of 20 ng cDNA, 2.5 μM of each forward and reverse primer, and 7.5 μl of 2x Power SYBR Green in 15 μl. The primers used were: Stat1 forward 5’-TAC GGA AAA GCA AGC GTA ATC T-3’ and reverse 5’-TGC ACA TGA CTT GAT CCT TCA C-3’; Socs3 forward 5’-ATG GTC ACC CAC AGC AAG TTT-3’ and reverse 5’-TCC AGT AGA ATC CGC TCT CCT-3’; Gapdh forward 5’-GTG TGA ACG GAT TTG GCC GTA-3’ and reverse 5’-GGA GTC ATA CTG GAA CAT GTA G-3’. The ΔΔCt method (81) was used for quantitating relative gene expression.

### High throughput sequencing and analysis

DNA extracted from tumors induced by TBLV-WT and TBLV-SD at the higher dose (3 tumors each) in wild-type and *Aicda^-/-^* B6 mice was used for PCR with primers *env*7254(+) (5’-ATC GCC TTT AAG AAG GAC GCC TTC T-3’) and LTR9604(-) (5’-GGA AAC CAC TTG TCT CAC ATC-3’), resulting in a 2.3 kb amplicon of the envelope-LTR junction region. Reactions were performed with JumpStart RED Accutaq LA polymerase (Sigma-Aldrich) in the supplied buffer, 1 µg of tumor DNA, 50 pmol of each primer, and 0.5 mM deoxynucleoside triphosphates in 20 µl. PCR parameters were: 94°C for 1 min, then 10 cycles at 94°C for 10 sec, 53°C for 30 sec, and 68°C for 2 min followed by 25 cycles of 95°C for 15 sec, 50°C for 30 sec, and 68°C for 2 min with a final incubation at 68°C for 7 min. Sixteen independent reactions for each tumor sample were pooled, and DNA was sheared to a size of approximately 400 bp using a Covaris ultrasonicator at the UT Austin Genomic Sequencing and Analysis Facility (GSAF). Fragmented DNA was end repaired and used for dA-tailing and ligation with Illumina adapters (unique to each tumor sample). Adapter-ligated DNA was amplified using these PCR parameters: 98°C for 30 sec, 6 cycles at 98°C for 10 sec, 65°C for 30 sec, and 72°C for 30 sec with a final incubation at 72°C for 5 min. Library preparations were submitted for sequencing on the MiSeq platform at GSAF.

For the high-throughput sequence analysis of TBLV proviruses from the tumors induced by low-dose infection, the same PCR fragment used for Sanger sequencing was purified using a Zymoclean^TM^ Gel DNA Recovery Kit (Zymo Research, cat# D4002). DNA (∼1 µg) was prepared for Nanopore long amplicon sequencing by the University of Wisconsin Biotechnology Center. After eliminating likely PCR duplicates and quantifying the number of reads aligned with each base at each position relative to the reference sequence, alternative base frequencies were dichotomized using a cutoff of 3%. For each pair (reference base to alternate base), the dichotomized alternate calls were modeled as a function of sample group (TBLV-SD versus TBLV-WT in either the wild-type or AID-knockout B6 mice). A mixed-effects logistic regression approach was used, incorporating a random effect that accounted for sample variation within the group.

### RNA extraction and RNA-seq analysis

Frozen thymic tumor tissue was ground to a fine powder with a mortar and pestle and lysed in TRI Reagent^TM^ Solution (ThermoFisher Scientific, cat#AM9738) using a microtube-sized Dounce homogenizer. Total RNA was prepared using the Direct-zol RNA miniprep kit (Zymo Research #R2050) following the manufacturer’s protocol. Total RNA (1.25 µg) was used for RNA Tag-Seq analysis at the UT Austin GSAF. Genes with significant differential expression (p<0.05) in the TBLV-SD-induced tumors were compared to those induced by TBLV-WT in the *Aicda*^-/-^ background and subjected to volcano plots and PANTHER pathway analysis (www.pantherdb.org).

### Splenocyte isolation and treatments

Spleens were removed aseptically and crushed to make a single-cell suspension in 5 ml of a solution of PBS plus 1% fetal bovine serum (FBS). Cells were subjected to centrifugation for 7 min at 335 Xg and resuspended in 3 ml of red blood lysis solution (9 parts 0.15 M NH_4_Cl plus 1 part 0.15 M Tris-HCl, pH 7.6). Following a 5-min incubation at room temperature, 20 ml of PBS/FBS solution was added, and the cells were pelleted again. The pellet was resuspended in 4 ml of PBS-FBS solution, filtered through a 40 µM cell strainer, and counted for viable cells using trypan blue.

For some experiments, 1 x 10^7^ splenocytes were added to 100mm Petri dishes containing 15 ml of stimulation media containing RPMI 1640 (Sigma R8758), antibiotic-antimycotic solution (Gibco 15240062), 1 mM sodium pyruvate (Sigma P5280), 10% FBS, 50 µM 2-mercaptoethanol, 5 ng/ml IL-4 (Sigma I1020), and 20 µg/ml LPS (Sigma L-2630). In other experiments, 3 x 10^7^ splenocytes were added to 100mm Petri dishes containing 15 ml of complete RPMI media (RPMI 1640, 10% FBS, 100U/ml penicillin, 100 µg/ml streptomycin, 2 mM L-glutamine, 50 µg/ml gentamycin sulfate) supplemented with 4 to 5 µg/ml ConA (Thermo Scientific AAJ61221MC). To determine the effects of type I interferons, 1 x 10^7^ cells were added to 100mm Petri dishes containing 15 ml of complete RPMI media plus 200 to 1000 U/ml IFN-β (R&D Systems 8234MB010). All cells were grown in a 37⁰C incubator with 7.5% CO_2_ for 5 h to 4 days (as indicated) prior to cell extract preparation.

### Cell lysate preparation and Western blotting

Whole cell extracts were prepared by adding either one volume of 2X-SDS-loading buffer [250 mM Tris-HCl, pH 6.8, 20% glycerol, 2% sodium dodecyl sulfate (SDS), 5% β-mercaptoethanol, 0.1% bromophenol blue] to cells in one volume of phosphate-buffered saline (PBS) or one volume of 6X-SDS loading buffer (0.35 M Tris-HCl, pH 6.8, 10% SDS, 36% glycerol, 0.6 M DTT, 0.012% bromophenol blue) to cells in five volumes of PBS. Samples were boiled for 5 min for protein denaturation. Protein concentrations were determined by Bradford assay (Bio-Rad Protein Assay Dye Reagent Concentrate, cat#5000006). The same amounts of whole cell lysates (20-25 µg) were loaded on 12% polyacrylamide gels and transferred onto nitrocellulose membranes (Cytiva Amersham™ Protran™ Supported NC Nitrocellulose Membranes, cat#10600037). After blocking with Intercept Blocking Buffer (LI-COR, cat#927-60001), blots were incubated with the primary antibody overnight at 4°C. The secondary antibody was incubated with membranes for 1.5 h at room temperature. After each antibody incubation, blots were washed three times for 5 min at room temperature in TBS-T (0.1% Tween-20, 20 mM Tris, 136 mM NaCl, pH 7.6). Bands were detected by LI-COR scanning or chemiluminescence (Amersham cat# RPN2232).

Antibodies to AID, p-Stat1(Y701), and Gapdh were obtained from Cell Signaling Technology. The mA3 antibody was kindly provided by Drs. Leonard Evans and Stefano Boi (NIAID Rocky Mountain Laboratories) (64, 69). Horseradish peroxidase-linked secondary antibodies were obtained from Cell Signaling Technology, whereas IRDye-linked secondary antibodies were obtained from LI-COR Biosciences.

### Statistical analysis and reproducibility

Results of Kaplan-Meier plots were analyzed for significance by Mantel-Cox log-rank tests. Statistical differences in the numbers of mutations/clone were evaluated by non-parametric Mann-Whitney tests. A p-value of <0.05 was considered to be significant. All experiments were repeated at least twice with similar results.

## ACKNOWLEDGMENTS

This work was funded by NIH grant R01 AI131660. We thank the Nussenzweig laboratory for kindly providing the *Aicda*^-/-^ mice on the B6 background. The generosity of Dr. Leonard (Pug) Evan’s laboratory in providing the mA3-specific antibody prior to his untimely death was greatly appreciated. We also valued the advice of Dr. Stefano Boi.

## REFERENCES

1. Temin HM. 1964. THE PARTICIPATION OF DNA IN ROUS SARCOMA VIRUS PRODUCTION. Virology 23:486–494.

2. Harris RS, Dudley JP. 2015. APOBECs and virus restriction. Virology 479–480:131–145.

3. Strebel K. 2013. HIV accessory proteins versus host restriction factors. Curr Opin Virol 3:692– 699.

4. Xu WK, Byun H, Dudley JP. 2020. The Role of APOBECs in Viral Replication. Microorganisms 8.

5. Dudley JP. 2023. APOBECs: Our fickle friends? PLOS Pathogens 19:e1011364.

6. Okeoma CM, Lovsin N, Peterlin BM, Ross SR. 2007. APOBEC3 inhibits mouse mammary tumour virus replication in vivo. Nature 445:927–930.

7. Derse D, Hill SA, Princler G, Lloyd P, Heidecker G. 2007. Resistance of human T cell leukemia virus type 1 to APOBEC3G restriction is mediated by elements in nucleocapsid. Proc Natl Acad Sci U S A 104:2915–2920.

8. Kao S, Khan MA, Miyagi E, Plishka R, Buckler-White A, Strebel K. 2003. The human immunodeficiency virus type 1 Vif protein reduces intracellular expression and inhibits packaging of APOBEC3G (CEM15), a cellular inhibitor of virus infectivity. J Virol 77:11398– 11407.

9. Holmes RK, Koning FA, Bishop KN, Malim MH. 2007. APOBEC3F can inhibit the accumulation of HIV-1 reverse transcription products in the absence of hypermutation. Comparisons with APOBEC3G. J Biol Chem 282:2587–2595.

10. Iwatani Y, Chan DSB, Wang F, Maynard KS, Sugiura W, Gronenborn AM, Rouzina I, Williams MC, Musier-Forsyth K, Levin JG. 2007. Deaminase-independent inhibition of HIV-1 reverse transcription by APOBEC3G. Nucleic Acids Res 35:7096–7108.

11. Harris RS, Bishop KN, Sheehy AM, Craig HM, Petersen-Mahrt SK, Watt IN, Neuberger MS, Malim MH. 2003. DNA deamination mediates innate immunity to retroviral infection. Cell 113:803–809.

12. Mangeat B, Turelli P, Caron G, Friedli M, Perrin L, Trono D. 2003. Broad antiretroviral defence by human APOBEC3G through lethal editing of nascent reverse transcripts. Nature 424:99– 103.

13. Zhang H, Yang B, Pomerantz RJ, Zhang C, Arunachalam SC, Gao L. 2003. The cytidine deaminase CEM15 induces hypermutation in newly synthesized HIV-1 DNA. Nature 424:94– 98.

14. Mehle A, Strack B, Ancuta P, Zhang C, McPike M, Gabuzda D. 2004. Vif overcomes the innate antiviral activity of APOBEC3G by promoting its degradation in the ubiquitin-proteasome pathway. J Biol Chem 279:7792–7798.

15. Sheehy AM, Gaddis NC, Malim MH. 2003. The antiretroviral enzyme APOBEC3G is degraded by the proteasome in response to HIV-1 Vif. Nat Med 9:1404–1407.

16. Stavrou S, Ross SR. 2015. APOBEC3 Proteins in Viral Immunity. J Immunol 195:4565–4570.

17. Hagen B, Kraase M, Indikova I, Indik S. 2019. A high rate of polymerization during synthesis of mouse mammary tumor virus DNA alleviates hypermutation by APOBEC3 proteins. PLoS Pathog 15:e1007533.

18. Singh GB, Byun H, Ali AF, Medina F, Wylie D, Shivram H, Nash AK, Lozano MM, Dudley JP. 2019. A Protein Antagonist of Activation-Induced Cytidine Deaminase Encoded by a Complex Mouse Retrovirus. mBio 10.

19. Indik S, Günzburg WH, Salmons B, Rouault F. 2005. A novel, mouse mammary tumor virus encoded protein with Rev-like properties. Virology 337:1–6.

20. Mertz JA, Simper MS, Lozano MM, Payne SM, Dudley JP. 2005. Mouse mammary tumor virus encodes a self-regulatory RNA export protein and is a complex retrovirus. J Virol 79:14737–14747.

21. Byun H, Das P, Yu H, Aleman A, Lozano MM, Matouschek A, Dudley JP. 2017. Mouse Mammary Tumor Virus Signal Peptide Uses a Novel p97-Dependent and Derlin-Independent Retrotranslocation Mechanism To Escape Proteasomal Degradation. MBio 8.

22. Byun H, Halani N, Gou Y, Nash AK, Lozano MM, Dudley JP. 2012. Requirements for mouse mammary tumor virus Rem signal peptide processing and function. J Virol 86:214–225.

23. Dultz E, Hildenbeutel M, Martoglio B, Hochman J, Dobberstein B, Kapp K. 2008. The signal peptide of the mouse mammary tumor virus Rem protein is released from the endoplasmic reticulum membrane and accumulates in nucleoli. J Biol Chem 283:9966–9976.

24. Byun H, Halani N, Mertz JA, Ali AF, Lozano MM, Dudley JP. 2010. Retroviral Rem protein requires processing by signal peptidase and retrotranslocation for nuclear function. Proc Natl Acad Sci USA 107:12287–12292.

25. Das P, Xu WK, Gautam AKS, Lozano MM, Dudley JP. 2022. A Retrotranslocation Assay That Predicts Defective VCP/p97-Mediated Trafficking of a Retroviral Signal Peptide. mBio e0295321.

26. Xu WK, Gou Y, Lozano MM, Dudley JP. 2021. Unconventional p97/VCP-Mediated Endoplasmic Reticulum-to-Endosome Trafficking of a Retroviral Protein. J Virol 95:e0053121.

27. Beutner U, Kraus E, Kitamura D, Rajewsky K, Huber BT. 1994. B cells are essential for murine mammary tumor virus transmission, but not for presentation of endogenous superantigens. J Exp Med 179:1457–1466.

28. Ardavín C, Martín P, Ferrero I, Azcoitia I, Anjuère F, Diggelmann H, Luthi F, Luther S, Acha-Orbea H. 1999. B cell response after MMTV infection: extrafollicular plasmablasts represent the main infected population and can transmit viral infection. J Immunol 162:2538–2545.

29. Golovkina TV, Dudley JP, Jaffe AB, Ross SR. 1995. Mouse mammary tumor viruses with functional superantigen genes are selected during in vivo infection. Proc Natl Acad Sci USA 92:4828–4832.

30. Golovkina TV, Dudley JP, Ross SR. 1998. B and T cells are required for mouse mammary tumor virus spread within the mammary gland. J Immunol 161:2375–2382.

31. Li J, Hakata Y, Takeda E, Liu Q, Iwatani Y, Kozak CA, Miyazawa M. 2012. Two Genetic Determinants Acquired Late in Mus Evolution Regulate the Inclusion of Exon 5, which Alters Mouse APOBEC3 Translation Efficiency. PLOS Pathogens 8:e1002478.

32. Okeoma CM, Huegel AL, Lingappa J, Feldman MD, Ross SR. 2010. APOBEC3 proteins expressed in mammary epithelial cells are packaged into retroviruses and can restrict transmission of milk-borne virions. Cell Host Microbe 8:534–543.

33. Muramatsu M, Sankaranand VS, Anant S, Sugai M, Kinoshita K, Davidson NO, Honjo T. 1999. Specific expression of activation-induced cytidine deaminase (AID), a novel member of the RNA-editing deaminase family in germinal center B cells. J Biol Chem 274:18470–18476.

34. Pacholczyk G, Suhag R, Mazurek M, Dederscheck SM, Koni PA. 2008. Generation of C57BL/6 knockout mice using C3H × BALB/c blastocysts. BioTechniques 44:413–416.

35. Watanabe H, Numata K, Ito T, Takagi K, Matsukawa A. 2004. INNATE IMMUNE RESPONSE IN TH1- AND TH2-DOMINANT MOUSE STRAINS. Shock 22:460.

36. Ball JK, Diggelmann H, Dekaban GA, Grossi GF, Semmler R, Waight PA, Fletcher RF. 1988. Alterations in the U3 region of the long terminal repeat of an infectious thymotropic type B retrovirus. J Virol 62:2985–2993.

37. Bhadra S, Lozano MM, Dudley JP. 2005. Conversion of mouse mammary tumor virus to a lymphomagenic virus. J Virol 79:12592–12596.

38. Mustafa F, Bhadra S, Johnston D, Lozano M, Dudley JP. 2003. The type B leukemogenic virus truncated superantigen is dispensable for T-cell lymphomagenesis. J Virol 77:3866– 3870.

39. Held W, Waanders GA, Shakhov AN, Scarpellino L, Acha-Orbea H, MacDonald HR. 1993. Superantigen-induced immune stimulation amplifies mouse mammary tumor virus infection and allows virus transmission. Cell 74:529–540.

40. Golovkina TV, Chervonsky A, Dudley JP, Ross SR. 1992. Transgenic mouse mammary tumor virus superantigen expression prevents viral infection. Cell 69:637–645.

41. Muramatsu M, Kinoshita K, Fagarasan S, Yamada S, Shinkai Y, Honjo T. 2000. Class switch recombination and hypermutation require activation-induced cytidine deaminase (AID), a potential RNA editing enzyme. Cell 102:553–563.

42. Wrona TJ, Lozano M, Binhazim AA, Dudley JP. 1998. Mutational and functional analysis of the C-terminal region of the C3H mouse mammary tumor virus superantigen. J Virol 72:4746– 4755.

43. Kitamura D, Roes J, Kühn R, Rajewsky K. 1991. A B cell-deficient mouse by targeted disruption of the membrane exon of the immunoglobulin mu chain gene. Nature 350:423– 426.

44. Ghosh S, Hoselton SA, Schuh JM. 2012. Mu-deficient mice possess B1 cells and produce IgG and IgE, but not IgA, following systemic sensitization and inhalational challenge in a fungal asthma model. J Immunol 189:1322–1329.

45. Brandt-Carlson C, Butel JS, Wheeler D. 1993. Phylogenetic and Structural Analyses of MMTV LTR ORF Sequences of Exogenous and Endogenous Origins. Virology 193:171–185.

46. Kozak C, Peters G, Pauley R, Morris V, Michalides R, Dudley J, Green M, Davisson M, Prakash O, Vaidya A. 1987. A standardized nomenclature for endogenous mouse mammary tumor viruses. J Virol 61:1651–1654.

47. Chen J, MacCarthy T. 2017. The preferred nucleotide contexts of the AID/APOBEC cytidine deaminases have differential effects when mutating retrotransposon and virus sequences compared to host genes. PLoS Comput Biol 13:e1005471.

48. Pham P, Bransteitter R, Petruska J, Goodman MF. 2003. Processive AID-catalysed cytosine deamination on single-stranded DNA simulates somatic hypermutation. Nature 424:103–107.

49. Pham P, Afif SA, Shimoda M, Maeda K, Sakaguchi N, Pedersen LC, Goodman MF. 2017. Activation-induced deoxycytidine deaminase: structural basis favoring WRC hot motif specificities unique among APOBEC family members. DNA Repair (Amst) 54:8–12.

50. MacMillan AL, Kohli RM, Ross SR. 2013. APOBEC3 inhibition of mouse mammary tumor virus infection: the role of cytidine deamination versus inhibition of reverse transcription. J Virol 87:4808–4817.

51. Halemano K, Guo K, Heilman KJ, Barrett BS, Smith DS, Hasenkrug KJ, Santiago ML. 2014. Immunoglobulin somatic hypermutation by APOBEC3/Rfv3 during retroviral infection. Proc Natl Acad Sci USA 111:7759–7764.

52. Mertz JA, Mustafa F, Meyers S, Dudley JP. 2001. Type B leukemogenic virus has a T-cell-specific enhancer that binds AML-1. J Virol 75:2174–2184.

53. Broussard DR, Mertz JA, Lozano M, Dudley JP. 2002. Selection for c-myc integration sites in polyclonal T-cell lymphomas. J Virol 76:2087–2099.

54. Pucillo C, Cepeda R, Hodes RJ. 1993. Expression of a MHC class II transgene determines both superantigenicity and susceptibility to mammary tumor virus infection. J Exp Med 178:1441–1445.

55. Ferreira BL, Ferreira ÉR, de Brito MV, Salu BR, Oliva MLV, Mortara RA, Orikaza CM. 2018. BALB/c and C57BL/6 Mice Cytokine Responses to Trypanosoma cruzi Infection Are Independent of Parasite Strain Infectivity. Frontiers in Microbiology 9.

56. Purdy A, Case L, Duvall M, Overstrom-Coleman M, Monnier N, Chervonsky A, Golovkina T. 2003. Unique resistance of I/LnJ mice to a retrovirus is due to sustained interferon gamma-dependent production of virus-neutralizing antibodies. J Exp Med 197:233–243.

57. Nusse R, Varmus HE. 1982. Many tumors induced by the mouse mammary tumor virus contain a provirus integrated in the same region of the host genome. Cell 31:99–109.

58. Peters G, Brookes S, Smith R, Dickson C. 1983. Tumorigenesis by mouse mammary tumor virus: evidence for a common region for provirus integration in mammary tumors. Cell 33:369– 377.

59. Rajan L, Broussard D, Lozano M, Lee CG, Kozak CA, Dudley JP. 2000. The c-myc locus is a common integration site in type B retrovirus-induced T-cell lymphomas. J Virol 74:2466–2471.

60. Broussard DR, Lozano MM, Dudley JP. 2004. Rorgamma (Rorc) is a common integration site in type B leukemogenic virus-induced T-cell lymphomas. J Virol 78:4943–4946.

61. Cantaert T, Schickel J-N, Bannock JM, Ng Y-S, Massad C, Oe T, Wu R, Lavoie A, Walter JE, Notarangelo LD, Al-Herz W, Kilic SS, Ochs HD, Nonoyama S, Durandy A, Meffre E. 2015. Activation-Induced Cytidine Deaminase Expression in Human B Cell Precursors Is Essential for Central B Cell Tolerance. Immunity 43:884–895.

62. Issaoui H, Ferrad M, Ghazzaui N, Lecardeur S, Cook-Moreau J, Boyer F, Denizot Y. 2020. Molecular analysis of γ1, γ3, and α class switch recombination junctions in APOBEC3-deficient mice using high-throughput sequencing. Cell Mol Immunol 17:418–420.

63. Langlois M-A, Kemmerich K, Rada C, Neuberger MS. 2009. The AKV murine leukemia virus is restricted and hypermutated by mouse APOBEC3. J Virol 83:11550–11559.

64. Boi S, Kolokithas A, Shepard J, Linwood R, Rosenke K, Van Dis E, Malik F, Evans LH. 2014. Incorporation of mouse APOBEC3 into murine leukemia virus virions decreases the activity and fidelity of reverse transcriptase. J Virol 88:7659–7662.

65. Stavrou S, Nitta T, Kotla S, Ha D, Nagashima K, Rein AR, Fan H, Ross SR. 2013. Murine leukemia virus glycosylated Gag blocks apolipoprotein B editing complex 3 and cytosolic sensor access to the reverse transcription complex. Proc Natl Acad Sci USA 110:9078–9083.

66. Prats AC, De Billy G, Wang P, Darlix JL. 1989. CUG initiation codon used for the synthesis of a cell surface antigen coded by the murine leukemia virus. J Mol Biol 205:363–372.

67. Low A, Okeoma CM, Lovsin N, de las Heras M, Taylor TH, Peterlin BM, Ross SR, Fan H. 2009. Enhanced replication and pathogenesis of Moloney murine leukemia virus in mice defective in the murine APOBEC3 gene. Virology 385:455–463.

68. Stavrou S, Zhao W, Blouch K, Ross SR. 2018. Deaminase-Dead Mouse APOBEC3 Is an In Vivo Retroviral Restriction Factor. J Virol 92:e00168–18.

69. Boi S, Ferrell ME, Zhao M, Hasenkrug KJ, Evans LH. 2018. Mouse APOBEC3 expression in NIH 3T3 cells mediates hypermutation of AKV murine leukemia virus. Virology 518:377–384.

70. Kawai T, Adachi O, Ogawa T, Takeda K, Akira S. 1999. Unresponsiveness of MyD88-Deficient Mice to Endotoxin. Immunity 11:115–122.

71. King LB, Corley RB. 1990. Lipopolysaccharide and dexamethasone induce mouse mammary tumor proviral gene expression and differentiation in B lymphocytes through distinct regulatory pathways. Mol Cell Biol 10:4211–4220.

72. Kane M, Case LK, Kopaskie K, Kozlova A, MacDearmid C, Chervonsky AV, Golovkina TV. 2011. Successful transmission of a retrovirus depends on the commensal microbiota. Science 334:245–249.

73. Kunimoto H, McKenney AS, Meydan C, Shank K, Nazir A, Rapaport F, Durham B, Garrett-Bakelman FE, Pronier E, Shih AH, Melnick A, Chaudhuri J, Levine RL. 2017. Aid is a key regulator of myeloid/erythroid differentiation and DNA methylation in hematopoietic stem/progenitor cells. Blood 129:1779–1790.

74. Su L, Shi Y-Y, Liu Z-Y, Gao S-J. 2022. Acute Myeloid Leukemia With CEBPA Mutations: Current Progress and Future Directions. Frontiers in Oncology 12.

75. Miller IJ, Bieker JJ. 1993. A novel, erythroid cell-specific murine transcription factor that binds to the CACCC element and is related to the Krüppel family of nuclear proteins. Mol Cell Biol 13:2776–2786.

76. Wierenga ATJ, Vellenga E, Schuringa JJ. 2010. Down-regulation of GATA1 uncouples STAT5-induced erythroid differentiation from stem/progenitor cell proliferation. Blood 115:4367–4376.

77. Yoshimura A, Ito M, Chikuma S, Akanuma T, Nakatsukasa H. 2018. Negative Regulation of Cytokine Signaling in Immunity. Cold Spring Harb Perspect Biol 10:a028571.

78. Golovkina TV, Jaffe AB, Ross SR. 1994. Coexpression of exogenous and endogenous mouse mammary tumor virus RNA in vivo results in viral recombination and broadens the virus host range. J Virol 68:5019–5026.

79. Mustafa F, Lozano M, Dudley JP. 2000. C3H mouse mammary tumor virus superantigen function requires a splice donor site in the envelope gene. J Virol 74:9431–9440.

80. Mertz JA, Lozano MM, Dudley JP. 2009. Rev and Rex proteins of human complex retroviruses function with the MMTV Rem-responsive element. Retrovirology 6:10.

81. Winer J, Jung CKS, Shackel I, Williams PM. 1999. Development and Validation of Real-Time Quantitative Reverse Transcriptase–Polymerase Chain Reaction for Monitoring Gene Expression in Cardiac Myocytesin Vitro. Analytical Biochemistry 270:41–49.

